# Optimal transport fate mapping resolves T cell differentiation dynamics across tissues

**DOI:** 10.64898/2026.02.24.707057

**Authors:** Alec L. Plotkin, Genevieve N. Mullins, William D. Green, Huitong Shi, H. Kay Chung, Natalie Stanley, J. Justin Milner

## Abstract

Immune responses evolve across time and tissues through coordinated programs of proliferation, differentiation, and migration, yet most single-cell measurements capture only static molecular snapshots. As a result, reconstructing how immune cells transition between alternative fates remains challenging, particularly for CD8 T cells, whose differentiation is highly dynamic and shaped by rapid expansion, contraction, and tissue trafficking. Here, we introduce an optimal transport-based fate mapping framework that reconstructs continuous CD8 T cell trajectories across time and tissues. Applied to longitudinal single-cell RNA-seq data from CD8 T cells responding to acute viral infection in mice, this approach accurately recapitulates population dynamics and resolves coherent effector and memory T cell differentiation trajectories. Extending the model to multiple tissues, we identify and experimentally validate temporally distinct waves of migration into the small intestine that give rise to divergent tissue-resident memory (Trm) fates, long-lived T cells crucial in immunosurveillance. By integrating optimal transport inference with time-resolved *in vivo* labeling, we demonstrate that CD52 marks recent tissue entrants and distinguishes them from Trm precursors. Finally, trajectory-guided analysis of transcription factor regulons reveals both shared and context-specific gene regulatory programs and identifies AP4 as a key regulator of circulating versus tissue-resident specification. These results establish optimal transport as a principled framework for reconstructing immune cell fate dynamics and provide a quantitative map of early events governing antiviral CD8 T cell differentiation across tissues.

## INTRODUCTION

Lymphocyte responses to infection represent one of the most dynamic differentiation programs in mammalian biology. This regulated process encompasses rapid clonal expansion, diversification into multiple functional states, migration across anatomical compartments, and long-term persistence as immunological memory^1–4^. From a computational perspective, these dynamics must be inferred from static molecular snapshots, even as individual cells continue to change state, proliferate, and undergo apoptosis over time^5–7^. These features underscore a central challenge in single-cell immune profiling: accurately reconstructing cell fates from independent and discrete timepoints in dynamic and migrating populations. Addressing this challenge is crucial for robust interpretation of single-cell datasets and for identifying molecular drivers that can be harnessed to steer immune cell differentiation toward therapeutically desirable states^8–10^.

Single-cell omics have enabled high-resolution atlases of lymphocyte states across time and tissues, revealing extensive heterogeneity in T cell responses^11–16^. However, translating these snapshots into mechanistic trajectories remains challenging because classical trajectory inference approaches^17–19^ are not optimized for lymphocyte dynamics. In particular, (i) robust proliferation and cell death violate steady-state and mass-conservation assumptions encoded in many embedding and pseudotime-based approaches; (ii) cell state cyclicity (e.g., dedifferentiation-like transitions or re-expression of “naive-like” programs) can confound trajectory directionality^20–22^; and (iii) migration between tissue compartments introduces mixtures of differentiation and transport that are difficult to disentangle without explicitly modeling population flow^23^. As a result, trajectory reconstructions that perform well in developmental systems can be unstable or ambiguous in T cell responses where unique kinetics, expansion patterns, and tissue trafficking critically influence the cell state composition of antigen-specific T cell populations.

Optimal transport (OT) offers a powerful and intuitive framework for modeling how lymphocytes change over time by directly linking cell populations sampled at different timepoints^24–26^. Rather than forcing cells into a single ordered trajectory, OT estimates how groups of cells at one timepoint give rise to groups of cells at the next, yielding a “transport map” that captures how cellular states evolve forward or backward in time. This perspective is particularly well suited for studying immune cell differentiation. OT explicitly models transitions between observed timepoints, avoiding reliance on global pseudotime orderings that can obscure branching or convergence in dynamic systems. In addition, OT can naturally account for changes in population size by allowing probability mass to expand or contract, enabling realistic modeling of clonal expansion and apoptosis, hallmarks of T cell responses during infection ^24^. Finally, because OT learns relationships between entire cell distributions rather than local transition directions, it is more robust in settings with cyclic differentiation paths, as observed in CD8 T cells, and readily applies to scenarios in which cells move between anatomical compartments, such as from circulation into tissues^20,27^.

Here, we apply OT-based trajectory inference to CD8 T cell differentiation during acute viral infection and extend it to a multi-compartment setting that jointly models circulating and tissue-resident T cell responses. The CD8 T cell response involves activation of pathogen-specific naive cells followed by rapid proliferation and diversification into effector and memory populations, tissue trafficking, and long-term persistence of heterogeneous memory states^2,4,28–30^. While many transcriptional regulators of effector and memory T cell responses have been described, it remains difficult to (i) quantify fate decisions and bifurcations at single-cell resolution across the full response, (ii) determine which features are stable markers of future fate (as opposed to transiently expressed features defined in single-timepoint data), and (iii) disentangle migration from differentiation in tissue-resident memory cells (Trm), where arrival to tissue is asynchronous and local microenvironments influence phenotypes^4^. Together, these gaps limit our ability to construct predictive, context-aware models of CD8 fate decisions and to identify regulators that can be harnessed for engineering durable CD8 T cell responses.

By applying OT fate mapping to longitudinal single-cell CD8 T cell datasets from the spleen and gut, we establish a unified “fate flow” framework that enables fluid interpretation of antiviral CD8 T cell population dynamics across time and tissue. We first demonstrate that OT-reconstructed programs faithfully recover expected infection kinetics, including expansion and contraction, and yield coherent flows between canonical circulating CD8 T cell populations. We then introduce a trajectory-centric integration strategy that clusters cells by transport-derived fate similarity, enabling identification of dominant differentiation programs and clarification of intermediate states that are often ambiguous with conventional static T cell state labels. Notably, we extend OT to a two-compartment framework to quantify the timing and directionality of egress into tissue and re-entry into circulation early in infection, enabling separation of migration-driven and differentiation-driven changes. This analysis reveals that Trm fate-specification is strongly shaped by time of arrival, linking early tissue entry to long-lived resident programs and later entry to more terminal effector-like programs. Finally, by establishing OT-derived lineage markers and transcription factor regulon activities across T cell compartments, we identify shared and context-specific regulatory modules and validate key predicted regulators experimentally.

This work establishes OT as a powerful framework for reconstructing lymphocyte fate transitions in systems shaped by proliferation, death, and migration, and provides a quantitative atlas of context-dependent CD8 T cell differentiation programs during active infection. While considering T cell states as discrete, static endpoints has been instrumental in providing insights into the molecular programs governing T cell fate^1,11,31,32^, immune responses are more accurately described as continuous, probabilistic shifts across a dynamic state space^33–36^. By explicitly modeling these transitions, our approach captures the dynamic nature of T cell differentiation and the intricacies of T cell specification and trafficking. Beyond acute infection, this framework is broadly applicable to diverse disease settings such as chronic infection^37,38^, autoimmunity^39^, and cancer, where T cell state transitions can be manipulated to enhance immunotherapy responses^10,40^.

## RESULTS

### Optimal transport reconstructs dynamic CD8 T fates during acute antiviral responses

To better resolve trajectories of antiviral CD8 T cell responses, we first outline the conceptual challenges of reconstructing differentiation paths from longitudinal single-cell data. CD8 T cell responses are inherently dynamic where individual cells change state in response to antigen and environmental cues, while the overall population simultaneously expands, contracts, and migrates across tissues. A useful abstraction is to consider this process as a probabilistic random walk through a transcriptional state space, in which cells move in directions shaped by underlying regulatory programs but are also influenced by stochastic variation. In parallel, CD8 T cells can divide or undergo apoptosis, leading to changes in population mass. Collectively, these processes describe how an initial distribution of cellular states evolves over time (Figure 1A). Although this framework captures key features of T cell biology, it highlights limitations of existing analysis approaches. Population-level models accurately describe expansion and contraction but do not capture the diversity of individual cell behaviors. In contrast, single-cell trajectory methods resolve heterogeneity but often ignore proliferation, death, and population turnover. These complementary limitations motivate the need for an approach that can integrate population dynamics with single-cell fate inference.

**Figure 1.**
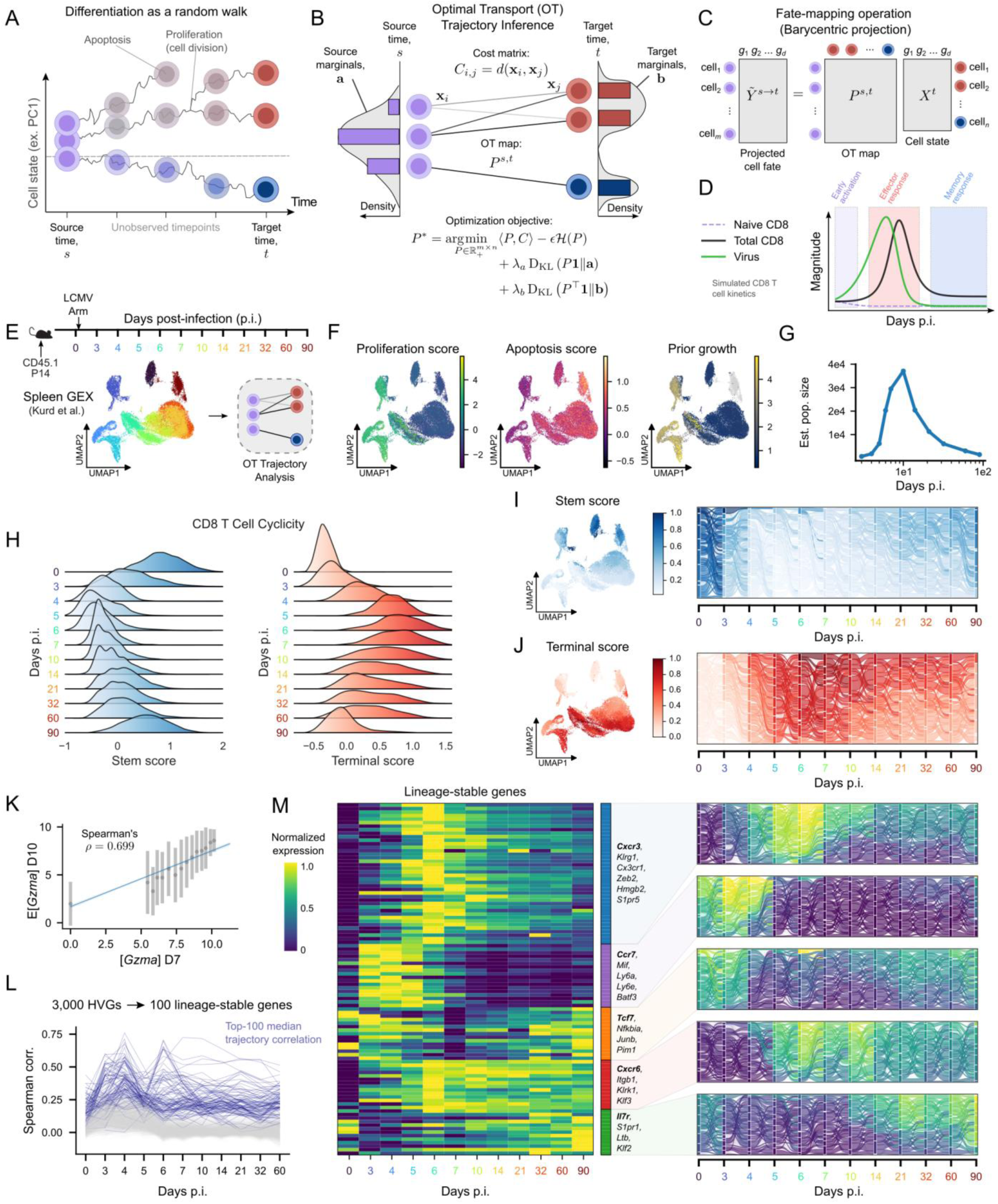
OT trajectory analysis reconstructs population dynamics of circulating CD8 T cells. **(A)** Abstraction of cellular differentiation as a random walk-through cell state space with birth (proliferation), death (apoptosis), and branching (differentiation) events. **(B)** Model illustrating the optimal transport (OT) objective, including marginal constraints and entropic regularization, which enable precise modeling of cellular differentiation. *a, b*: probability mass vectors for source and target marginals, respectively; *P*^*s,t*^: optimal transport map from timepoints *s* to *t*; *P*^∗^: optimal transport solution; *C*: transport cost matrix; *x*_*i*_, *x*_*j*_: cell state vectors; *d*(⋅,⋅): distance function; ℋ(⋅): Shannon entropy; ϵ: entropic regularization weight; *D*_*KL*_ (⋅ ║ ⋅): Kullback–Leibler divergence; λ_*a*_, λ_*b*_: unbalanced OT regularization weights. **(C)** Illustration of the fate-mapping operation, allowing OT to be used for fate prediction. **(D)** Population dynamics of CD8 T cell responses to acute infection, simulated using flexible fate decision model from Abadie et al.^43^ **(E)** Experimental summary of scRNAseq time course profiling of circulating P14 splenocytes in mice infected with LCMV Arm (from Kurd et al.^13^). UMAP dimensionality reduction of gene expression data shaded by timepoint. **(F)** UMAPs shaded by proliferation (left), apoptosis (middle), and prior growth rate scores (right) used to compute the input OT source marginals. **(G)** Estimated population size (from E), reconstructed from OT model. **(H)** Ridgeline plots displaying enrichment of stem (left) and terminal (right) gene set scores over time. **(I-J)** UMAPs (left) and Sankey diagrams (right) colored by supercell stem (I) and terminal (J) gene set scores. Flows in Sankey diagrams correspond to transport probabilities from OT maps. **(K)** Correlation between *Gzma* expression at day 7 and predicted *Gzma* expression at day 10 guided by OT mapping. **(L)** Traces of gene expression correlations induced by OT across time points. Top 100 genes with highest median correlations are outlined in blue. **(M)** Heatmap of normalized mean gene expression for the top 100 lineage-stable genes (left) and select Sankey diagrams (right) colored by supercell expression for representative lineage-stable genes from each module.

Motivated by this computational and conceptual disconnect, we utilized an OT framework (Figure 1B) to fit cell trajectories, defined as the chronological adoption of cell-states over time. OT is capable of modeling stochastic dynamics by probabilistically matching cells from a source distribution to their most likely descendants in a target distribution^24,25,41^. Importantly, this model allows us to implicitly account for proliferation and death by adjusting the amount of weight placed on each cell in the source distribution or “source marginals.” The output of OT between two timepoints is a transport map, which can be used to predict future or past cell states (Figure 1C). Unlike approaches that explicitly estimate local velocity fields, OT infers trajectories directly from distribution-to-distribution mappings, making it well-suited for systems with branching, convergence, and population turnover. These properties are especially well matched for modeling CD8 T cell responses to infection.

Antigen recognition through the T cell receptor triggers activation and rapid clonal expansion, followed by T cell trafficking through lymphoid organs to peripheral sites of inflammation. In acutely resolving infection such as lymphocytic choriomeningitis virus (LCMV) Armstrong (Arm), waning antigen levels leads to widespread effector cell apoptosis, while a subset of cells sustains pro-survival signaling and give rise to long-lived memory populations^42^. These patterns can be captured by coarse-grained models of CD8 T cell kinetics, which accurately reproduce population expansion and contraction over time^43,44^,45,46. We performed one such simulation using a flexible fate-decision model to illustrate these characteristic dynamics (Figure 1D), providing a quantitative baseline against which single-cell–resolved fate inference can be evaluated.

To investigate how single-cell heterogeneity evolves during the antiviral CD8 T cell response, we leveraged temporal OT via moscot^26^ in a longitudinal scRNA-seq dataset of antigen-specific “P14” CD8 T cells expressing a transgenic TCR recognizing the GP_33-41_ epitope of LCMV^12,13^. LCMV Arm induces an acutely resolving infection that elicits a robust, systemic CD8 T cell response, permitting analysis of differentiation dynamics across time and tissues. Uniform manifold approximation and projection (UMAP) visualization^47^ revealed that P14 cells sampled from days 0-90 post-infection (p.i.) clustered by timepoint as anticipated (Figure 1E). To estimate source marginals for OT, we computed proliferation and apoptosis gene set scores and combined them to derive a prior growth rate for each cell (Figure 1F). Consistent with expected response kinetics, gene set scores for proliferation peaked at days 4-5 p.i. whereas apoptosis scores increased gradually over time (Figure 1F, S1A). Using these growth-rate-derived source marginals, we estimated total CD8 T cell population sizes over time from the OT model (Figure 1G). Notably, these estimates recapitulated the expected phases of expansion and contraction and closely matched both the flexible fate-decision simulation (Figure 1D) and prior experimental measurements^45,46,48^, despite near-uniform cell sampling across timepoints. This finding demonstrated that OT inference accurately recovered population dynamics from snapshot data alone. We further performed multiple quality control analyses to confirm that OT inference was not driven by technical biases (Figures S1B–D).

We next assessed whether OT reconstructed key features of CD8 T cell differentiation, including anticipated patterns of stem-like and terminal differentiation programs that have been deeply investigated in response to infection^1^. Consistent with established CD8 T cell differentiation kinetics, terminal differentiation programs predominated during the effector phase of the response (∼days 5-10 p.i.), whereas cyclical stemness programs declined rapidly following activation and reemerged during memory formation (≥30 days p.i., Figure 1H). To assess whether OT captured these dynamics, we coarsened the data into timepoint-specific supercells and visualized OT-derived probability flows between them (Figure 1I–J). We use the term “fate flow” to denote the time-resolved redistribution of clonal or phenotypic mass between CD8 T cell fates, visualized as Sankey diagrams derived from our trajectory inference. Fate flows revealed streamlines that preferentially connected cells with similar stemness or terminal differentiation scores, highlighting continuity in differentiation that is obscured by static state labels. This phenotypic coherence supports that OT accurately reconstructs CD8 T cell fate dynamics during infection.

Finally, we used OT trajectories to identify stable gene expression programs along differentiation paths, rather than transiently expressed individual genes from specific cell states (as is typical in conventional static analyses). In dynamic immune responses, many commonly used lineage markers, such as *Klrg1*, show strong but short-lived expression patterns, which can complicate their interpretation in static single-cell analyses. Because OT explicitly links cells across time, it provides a way to determine whether a gene’s expression is maintained as cells progress along a particular differentiation trajectory. Here, we used OT transport maps to predict each cell’s expected gene expression at the subsequent timepoint and compared these predictions to observed expression (as with *Gzma* in Figure 1K). Genes whose expression closely matched their OT-based predictions were consistently propagated along inferred differentiation paths, indicating stable association with fate rather than transient expression (Figure 1M). Across all genes and adjacent timepoints, we identified the top 100 genes with the highest median Spearman correlation between observed and OT-projected expression (Figure 1L). Clustering these genes by their temporal patterns revealed distinct modules corresponding to different phases of the immune response (Figure 1M). Visualization of fate-flow modeling recovered well-established lineage markers (e.g., *Tcf7, Il7r, Ccr7*) and highlighted more nuanced dynamics for genes such as *Cxcr3* and *Cxcr6*. These analyses show that OT-based fate-flow reconstruction provides a practical way to distinguish stable lineage programs from transient state-dependent gene expression during CD8 T cell differentiation.

### Trajectory clustering exposes fluid differentiation paths obscured by static CD8 T cell nomenclature

Circulating antiviral CD8 T cell differentiation is traditionally organized into discrete cell states defined by phenotype, function, anatomical localization, and infection context. In this framework (Figure 2A), early effector CD8 T cells (Tee) arise from newly activated CD8 T cells by approximately day 4 post-infection and are thought to give rise to both terminal effector (Tte) and memory precursor (Tmp) populations^49,50^. As viral antigen is cleared, a subset of effector cells transitions into long-lived memory states, with Tmp cells predominantly giving rise to effector memory (Tem) and central memory (Tcm) populations, whereas a smaller fraction of terminal effector cells differentiates into terminal effector memory (Ttem) cells^50,51^. This state-based model has provided a useful and interpretable framework for linking molecular programs to immune function, and gene set-based annotation of our data recapitulated these canonical circulating CD8 T cell populations (Figure 2B). However, when viewed through the lens of fate flows, these static assignments (Figure 2B) or conventional clustering assignments (Figure S1E) highlight important limitations. Transitions between labeled states appear abruptly, obscuring the continuity of differentiation paths and making it difficult to resolve how individual effector populations relate quantitatively to downstream memory fates. This ambiguity is further reflected in population dynamics, where linking specific effector states to expanding or persisting memory pools remains challenging (Figure 2B, right). These observations suggest that while T cell subset nomenclature has been invaluable in our understanding of CD8 T cell biology, it incompletely captures the fluid, continuous structure of CD8 T cell differentiation over time.

**Figure 2.**
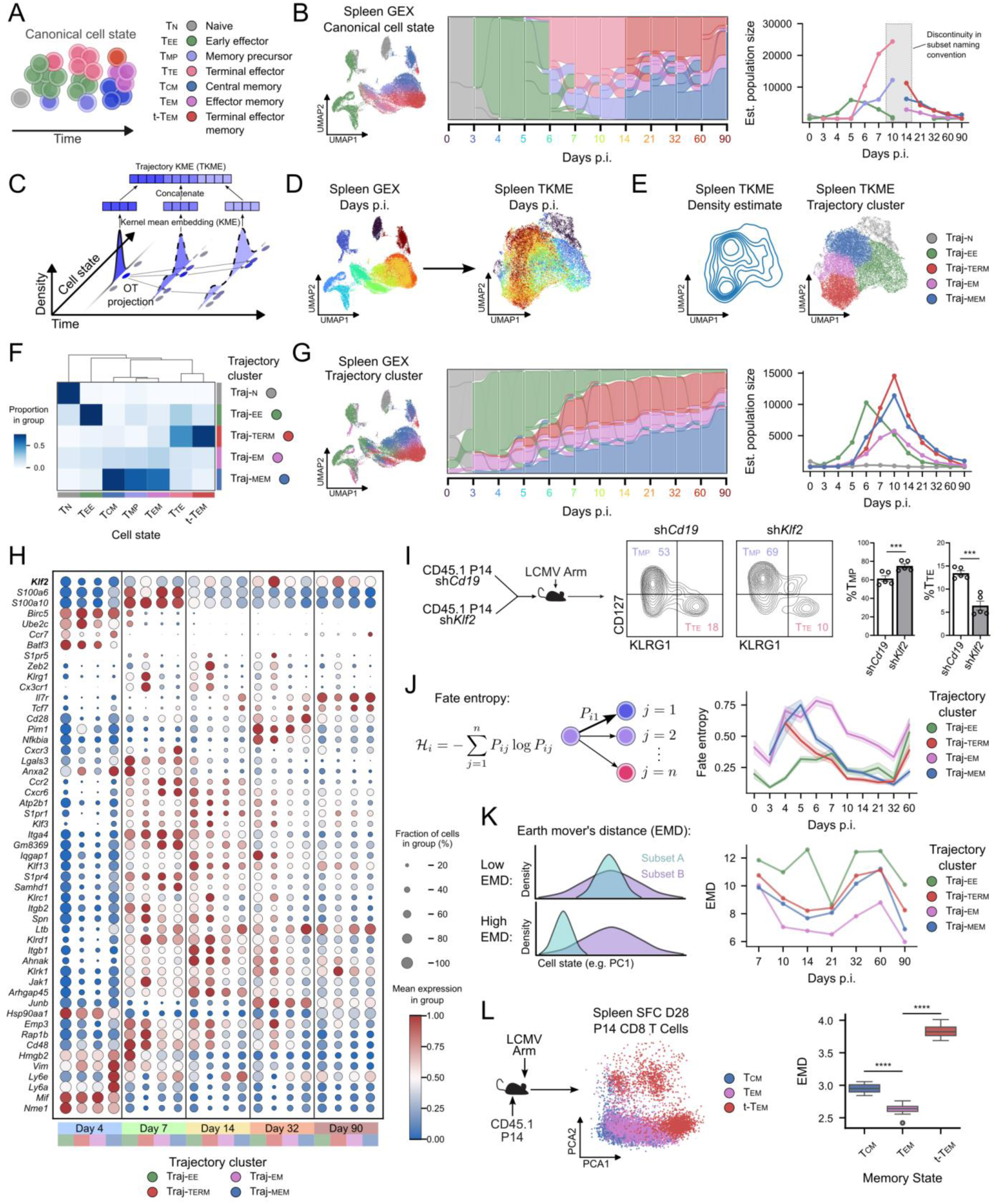
Trajectory clustering exposes fluid differentiation paths obscured by static CD8 T cell nomenclature. **(A)** Schematic representing canonical circulating CD8 T cell states (based on static nomenclature) in acute infection. **(B)** Canonical cell states projected in GEX UMAP (left), Sankey diagram (middle), and population kinetics reconstruction (right). **(C)** Schematic of OT-based trajectory integration approach. **(D)** UMAPs derived from gene expression (GEX; left) and trajectory kernel mean embeddings (TKME; right), shaded by infection timepoint. **(E)** Kernel density estimate (left) and leiden clusters (right) plotted in TKME UMAP space. **(F)** Confusion matrix displaying overlap between canonical cell states (columns) and trajectory clusters (rows), normalized by column. **(G)** Trajectory clusters projected in GEX UMAP space (left), Sankey diagram (middle), and population kinetics reconstruction (right). **(H)** Dotplot of gene expression for trajectory clusters at representative time points (days 4, 7, 14, 32, and 90 p.i.). Genes correspond to top lineage genes derived from Figure 1M, filtered to display genes encoding cell-surface proteins, secreted proteins, or transcription factors. **(I)** Congenic P14 cells transduced with sh*Cd19*-encoding retrovirus (control) or sh*Klf2*-enoding retrovirus were mixed 1:1 and transferred into mice infected with LCMV Arm. Representative flow cytometry plots and quantification are shown. Graphs show mean+/-SEM, each dot represents an individual mouse, students t test, n=5, *p<0.005. **(J)** Schematic (left) of fate entropy, and line plots (right) showing mean fate entropy for each cluster across time. Shaded regions represent 95% confidence intervals. **(K)** Schematic (left) of Earth Mover’s Distance (EMD) as a metric for distributional distance, and line plots (right) displaying EMDs of each trajectory cluster over time. **(L)** Congenic P14 cells were transferred into mice infected with LCMV Arm. On day 28 p.i. spectral flow cytometry analysis (SFC) was performed on splenocytes and PCA was performed. Boxplots (right) show EMDs of each cell state against all other cells, across 10 bootstraps of 600 cells each. 2-sided t-test, ****p<0.001.

To establish a more coherent computational model for fitting fluid differentiation paths, we developed an unsupervised approach that clusters cells based on the similarity of their OT-derived trajectories. We first constructed per-cell trajectory kernel mean embedding (TKME; Figure 2C) representations, and projected the cells using UMAP according to the computed trajectory embeddings. This allowed us to characterize cells based on their entire trajectories, rather than their transient states. We found that, in contrast to gene expression (GEX)-derived embeddings, the TKME embeddings effectively integrated cells across timepoints (Figure 2D). Within the TKME UMAP space, we observed distinct distributions which we hypothesized corresponded to separate CD8 T cell differentiation paths (Figure 2E, left). We then performed Leiden clustering in this space (Figure 2E, right) and labeled clusters based on key CD8 T cell gene set signatures reflecting effector and memory trajectories (Figure 2E, Figure S1F). TKME clustering enabled delineation of the following distinct trajectories: long-lived memory (Traj-mem), terminal fated (Traj-term), early effector (Traj-ee), and intermediate effector memory (Traj-em). We found that the trajectory clusters closely corresponded to the expected canonical cell states. For example, Traj-mem overlapped with Tcm, Tmp, and to a lesser extent Tem states, while Traj-term overlapped with both Tte and Ttem. Traj-em exhibited an intermediate distribution between stem-like memory cells and terminally fated populations (Figure 2F). Critically, visualization of fate flows (Figure 2G) revealed that trajectory clusters were coherent over time, and outperformed both gene set score annotation as well as clustering in GEX space with respect to trajectory consistency (Figures S1G).

Trajectory clustering provided temporal structure to the OT-derived CD8 T cell differentiation response (Figure 2G). We observed a wave of cells within Traj-ee that expands between days 0-4 p.i. followed by sharp population contraction on days 5-10 p.i. Further, we observed Traj-term emerged at days 5-6 p.i. (consistent with the emergence of Tte cells) and was maintained until approximately D60, consistent with Ttem contraction dynamics^50^. The Traj-MEM cells continued to expand over time, consistent with Tcm persisting late into the course of acute infection. Last, we observed that Traj-EM emerged early at D3 and remained relatively stable throughout the course of infection. In summary, fate flow reconstruction of T cell differentiation trajectories supported existing expectations of effector and memory T cell differentiation.

Analysis of gene expression within trajectory clusters provided an orthogonal and potentially more robust view of CD8 T cell differentiation by grouping cells according to shared fates rather than conventional clustering (e.g. static transcriptional similarity). Because trajectory clusters aggregate cells that participate in the same differentiation paths across time, gene expression patterns within these clusters are less influenced by transient activation or cell cycle effects and instead reflect stable lineage-associated programs. Consistent with established CD8 T cell biology, trajectory clusters displayed distinct temporal and path-dependent expression patterns (Figure 2H). For example, *Klf2* expression increased over time across all trajectories but was most pronounced within Traj-TERM beginning at D7, in agreement with its known role in regulating effector differentiation and migration^52,53^. Guided by trajectory-resolved expression patterns, we assessed a cell-intrinsic role of KLF2 in Tte differentiation during acute LCMV infection. Consistent with prior studies implicating KLF2 in shaping effector programs^11,52^, genetic silencing of *Klf2* impaired formation of Tte-like cells while increasing the frequency of Tmp-like cells on day 14-15 p.i. (Figure 2I). Similarly, *Zeb2*, a well-characterized driver of terminal differentiation in acute infection^11,52^, was selectively upregulated within Traj-TERM between days 7-32 p.i., further supporting the biological coherence of the inferred trajectories.

For most lineage-associated genes, Traj-EM exhibited expression levels that were intermediate between Traj-MEM and Traj-TERM (Figure 2H), consistent with the effector-memory nature of Tem cells that likely exist on a continuum between stem-like Tcm and terminally differentiated Ttem. Traj-em cells also showed higher uncertainty in fate assignment than other trajectories, as quantified by Shannon entropy of fate probabilities (Figure 2J), potentially reflecting their functional plasticity. Indeed, Tem cells have previously been shown to give rise to Tcm but also exhibit self-renewal properties to maintain the Tem pool, in contrast to Ttem that remain largely fate locked with limited self-renewal potential^50^. Accordingly, in the trajectory embedding space, Traj-em cells occupied an intermediate position between Traj-term and Traj-mem fates (Figure 2E). To quantitatively assess this intermediate positioning, we measured earth mover’s distance (EMD) between Traj-em and other trajectories over time. Consistent with an intermediate or transitional identity, the Traj-em trajectory exhibited the lowest EMD relative to other trajectories at later timepoints (Figure 2K), supporting the interpretation that these cells bridge terminal and memory fates. This relationship was also supported by high-dimensional spectral flow cytometry profiling of memory CD8 T cell populations on day 28 p.i., where Tem cells were interpolated between Tcm and Ttem populations (Figure 2L). Consistent with the trajectory-based analysis, Tem cells also exhibited the lowest EMD among circulating memory states in protein expression spectral flow cytometry space. Taken together, these results across transcriptomic and proteomic analyses show that trajectory-based fate flow reconstruction reliably recapitulates established features of CD8 T cells and their differentiation hierarchies, while offering a quantitative way to interpret intermediate states that are difficult to resolve using static classifications alone.

### Multi-compartment optimal transport reveals CD8 T cell migration dynamics

A hallmark of the CD8 T cell response is trafficking to sites of inflammation, where they can differentiate into specialized tissue-resident populations that control pathogen burden in infected tissues. Tissue-resident CD8 T cells are critical mediators of immunosurveillance, providing long-term sentinel protection at barrier sites of frequent pathogen exposure^54^. Although recent studies have begun to define precursors of tissue-infiltrating CD8 T cells and their differentiation into long-lived Trm^11,13,16^, the early signals that govern Trm fate specification remain incompletely understood. A central challenge is that CD8 T cells are recruited asynchronously from lymphoid tissues and circulation into infected non-lymphoid sites, making it difficult to reconstruct temporal differentiation paths from static scRNA-seq snapshots. We therefore sought to determine whether OT-based trajectory inference could reconstruct coherent fate flows between two anatomically distinct compartments, including lymphoid tissue harboring circulating CD8 T cells and non-lymphoid tissue containing resident cells.

LCMV Arm induces a systemic infection that involves multiple tissues sites, including robust and well-characterized CD8 T cell responses in the spleen and the small intestine intraepithelial lymphocyte (siIEL) compartment. Using longitudinal scRNA-seq measurements of antigen-specific P14 CD8 T cells from both the spleen and siIEL (Figure 3A, left), UMAP visualization revealed that cells from these compartments occupy distinct, largely non-overlapping transcriptional spaces, consistent with prior reports^12,13^. Previous work has shown that LCMV-specific CD8 T cells traffic into the siIEL within a discrete window approximately 3-7 days p.i.^23^ Outside this window, ingress and egress are constrained, providing a well-defined setting to evaluate OT-inferred cellular mixing between compartments.

**Figure 3:**
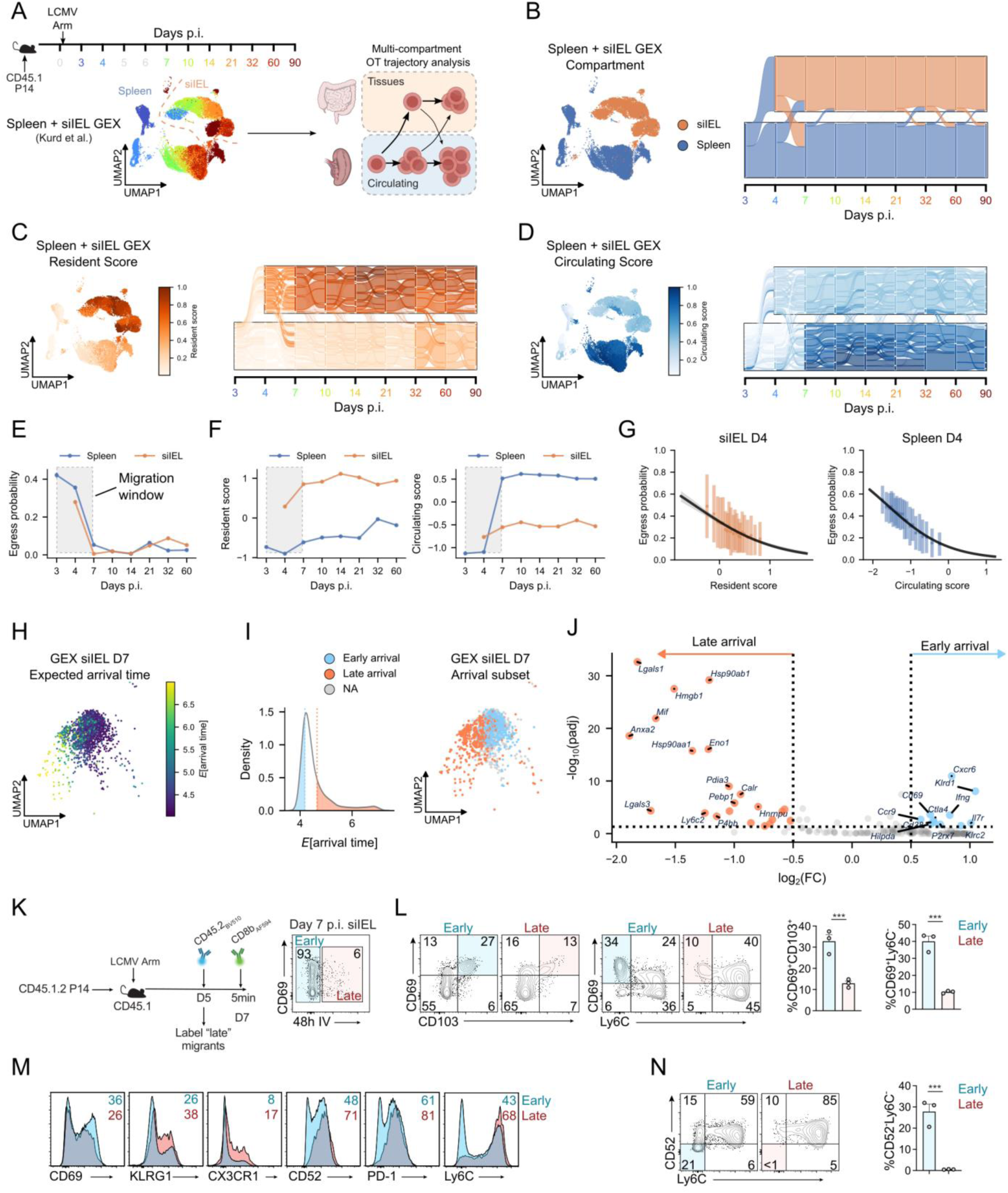
Multi-compartment optimal transport reveals CD8 T cell migration dynamics. **(A)** Summary of scRNAseq experiment profiling circulating and tissue-resident P14 cells (left, sequencing data from Kurd et al.). Cells were isolated from the spleen and small intestine intraepithelial compartment (siIEL). Days where splenocytes but not siIEL cells collected are greyed out on the time scale axis. UMAP represents combined spleen and siIEL GEX, shaded by day p.i. Cartoon (right) illustrating the intuition underlying the multi-compartment OT trajectory inference model. **(B)** UMAP (left) and Sankey diagram of cross-compartment trajectories (right) shaded by tissue compartment. **(C-D)** UMAP (left) and Sankey diagram (right) colored by tissue-resident (C) and circulating (D) gene set scores. **(E)** Line plots of tissue egress/migration probabilities from spleen (blue) and siIEL (orange) over time. Error bars represent mean probability +/-SEM. The shaded box corresponds to an experimentally determined migration window^23^ for siIEL cells. **(F)** Line plots of resident (left) and circulating (right) scores for spleen and siIEL over time with migration window highlighted in grey. **(G)** Correlation plots of circulating scores vs. migration probabilities at day 4 p.i. in spleen (left) and siIEL (right). Lines of best fit correspond to logistic regression models of egress probability versus resident (odds ratio = 0.287, P = 5.8e-10) or circulating (odds ratio = 0.289, P = 1.1e-18) scores. **(H)** UMAP of siIEL cells isolated at 7 day p.i., with colorbar representing predicted arrival times into the siIEL from circulation. **(I)** Histogram (left) of arrival times, with the top and bottom quartiles highlighted. UMAP (right) with cells from the bottom and top quartiles labeled as early and late arrivals, respectively. **(J)** Volcano plot of differentially expressed cell surface genes in early vs. late arrivals. FC: fold change; padj: Benjamini-Hochberg adjusted p-values. **(K)** Schematic (left) of intravenous (IV) labeling experiment to distinguish early vs. late arrivals (CD45.2 antibodies) and blood vs tissue localized (CD8b antibody) on day 7 p.i. Flow cytometry plot (right) displaying proportion of late migrants into the siIEL. **(L)** Flow plots and quantification of frequency of CD69^+^CD103^+^ and CD69^+^Ly6c^-^ cells in early (IV-) and late (IV+) arrivals to the siIEL (from K). **(M)** Frequency of cells expressing high levels of CD69, KLRG1, CX3CR1, CD52, PD-1, and Ly6c in early vs. late arrivals (from K). **(N)** Flow cytometry plots and quantification of CD52 and Ly6c expression in early and late arrivals to the siIEL (from K). Graphs show mean+/-SEM, n=3/group, one representative of two independent experiments, ****p<0.001 (L-N).

To approximate inter-compartmental CD8 T cell migration dynamics, we fit OT trajectories with day 3 p.i. P14 splenocytes as the initiating timepoint, resulting in a coarse model where it was assumed that T cells can belong to one of two compartments: circulating or tissue-resident, represented by CD8 T cells from the spleen or the siIEL, respectively (Figure 3A, right). This framework allowed us to estimate the probability of a given cell to change compartments from one timepoint to the next, visualized as fate flows between circulation and tissue (Figure 3B). As in the spleen-only analysis, we incorporated proliferation and apoptosis into a prior growth rate estimate and performed extensive quality control to ensure that the multi-compartment OT model was free of systematic biases (Figures S2A-C). In order to prevent artifacts from nonconstant compartment sample sizes over time, we subsampled at each timepoint to ensure that both compartments had equal numbers of cells.

The model predicted migration predominantly from the spleen into the intestine, consistent with established spleen-to-siIEL trafficking^23^. Most P14 cells were predicted to arrive in the siIEL by day 4 post-infection, with a smaller fraction arriving between days 4–7 p.i. (Figure 3B). A modest population of siIEL cells was also predicted to migrate from the intestine back into circulation during this interval. After day 7, both ingress and egress probabilities sharply declined (Figure 3B), in agreement with prior studies of CD8 T cell trafficking to the intestine during acute infection^23^.

We and others have shown that a core tissue-residency gene expression program is rapidly engaged upon T cell infiltration into non-lymphoid sites, which facilitates T cell lodging and differentiation into bona fide Trm^55,56^. Notably, the OT-predicted migration window coincided with dynamic shifts in circulating and resident gene expression programs (Figure 3C-E). P14 cells localized to the siIEL rapidly upregulated residency-associated genes, whereas splenic cells reinforced circulating gene expression programs, underscoring the plasticity of these programs during the migration window (Figure 3C-E). Fate-flow analysis further showed that cells with high residency gene set scores tended to remain in the siIEL (Figure 3C,F), while cells with high circulating scores remained confined to circulation (Figure 3D,F). Within each tissue, stronger compartment-associated gene signatures were inversely correlated with egress probability at day 4 post-infection (Figure 3G), indicating that the degree of tissue adaptation is a key determinant of CD8 T cell migration dynamics.

To investigate early molecular events guiding Trm specification, we analyzed genes associated with predicted time of arrival in the siIEL. Using the multi-compartment OT model, we estimated arrival times for siIEL-localized cells at day 7 p.i. (Figure 3H), revealing a distribution skewed toward early arrivals (Figure 3I, left). Defining the top and bottom quartiles as early and late arrivals, respectively, we found that these populations occupied distinct regions of GEX space (Figure 3I, right). Differential expression analysis revealed that early arrivals preferentially expressed residency- and memory-associated genes (*Cd69, P2rx7, Cxcr6, Ccr9, Il7r*), whereas late arrivals upregulated effector- and circulation-associated genes (*Lgals1, Lgals3, Mif, Ly6c2*) (Figure 3J). These findings suggest that asynchronous arrival contributes to phenotypic heterogeneity among early siIEL cells and underscores the importance of explicitly modeling migration dynamics in immune fate-mapping analyses.

To experimentally validate these computational predictions, we performed time-resolved intravascular antibody labeling of donor CD45.1/CD45.2 P14 CD8 T cells in congenic CD45.1/CD45.1 LCMV Arm-infected mice (Figure 3K). CD45.2 antibody conjugated to a fluorophore suitable for longitudinal labeling^57^ was administered intravenously 48 hours prior to tissue harvest at day 7 p.i., selectively labeling circulating but not tissue-localized cells (Figures 3K). Antibody-labeled P14 cells present in the siIEL on day 7 p.i. represented “late arrival” cells localized in the siIEL for less than 48h, distinct from unlabeled “early arrival” cells presumably lodged in the siIEL for >48h. Consistent with OT predictions, >90% of siIEL P14 cells were comprised of early arrivals on day 7 p.i. (Figure 3K).

Early-arrival P14 cells exhibited a more mature Trm-like profile, with a higher frequency of CD69^+^CD103^+^ cells and CD69^+^Ly6C^-^ cells, presumably due to a longer period of exposure to Trm-inducing cues in the siIEL (Figure 3L). Conversely, late arrivals expressed higher levels of KLRG1, CX3CR1, CD52, and PD-1, consistent with an effector-like, circulating phenotype (Figure 3M). These experimentally observed expression patterns closely matched OT-guided predictions (Figure 3J). Further, we found that, despite harvesting siIEL cells at a single infection timepoint, CD52 and Ly6C sharply discriminated asynchronous infiltration of early (CD52^-^Ly6C^-^) from late (CD52^+^Ly6C^+^) arriving cells *in vivo* (Figure 3N). These results validate the OT-based migration model, highlight CD52 as a novel marker for distinguishing distinct tissue-resident cell states, and demonstrate that time of tissue entry imprints phenotypic differences in siIEL CD8 T cells.

### Migration kinetics influence differentiation of Trm precursors

Differences between early and late arrivals in the siIEL could arise from intrinsic differences in the pace of differentiation after entry or changes in the local siIEL microenvironment, but they may also reflect discrete differences in the circulating precursor populations that seed the gut at different timepoints. We sought to leverage our OT-based fluid model of differentiation to glean insight into the degree to which tissue-resident phenotypes are imprinted in circulating cells. To address this, we overlaid spleen-derived trajectories onto our multi-compartment integration model to resolve the fate-flows comprising early (∼days 3-4 p.i.) and late (∼days 4-7 p.i.) waves of tissue entry (Figure 4A).

**Figure 4:**
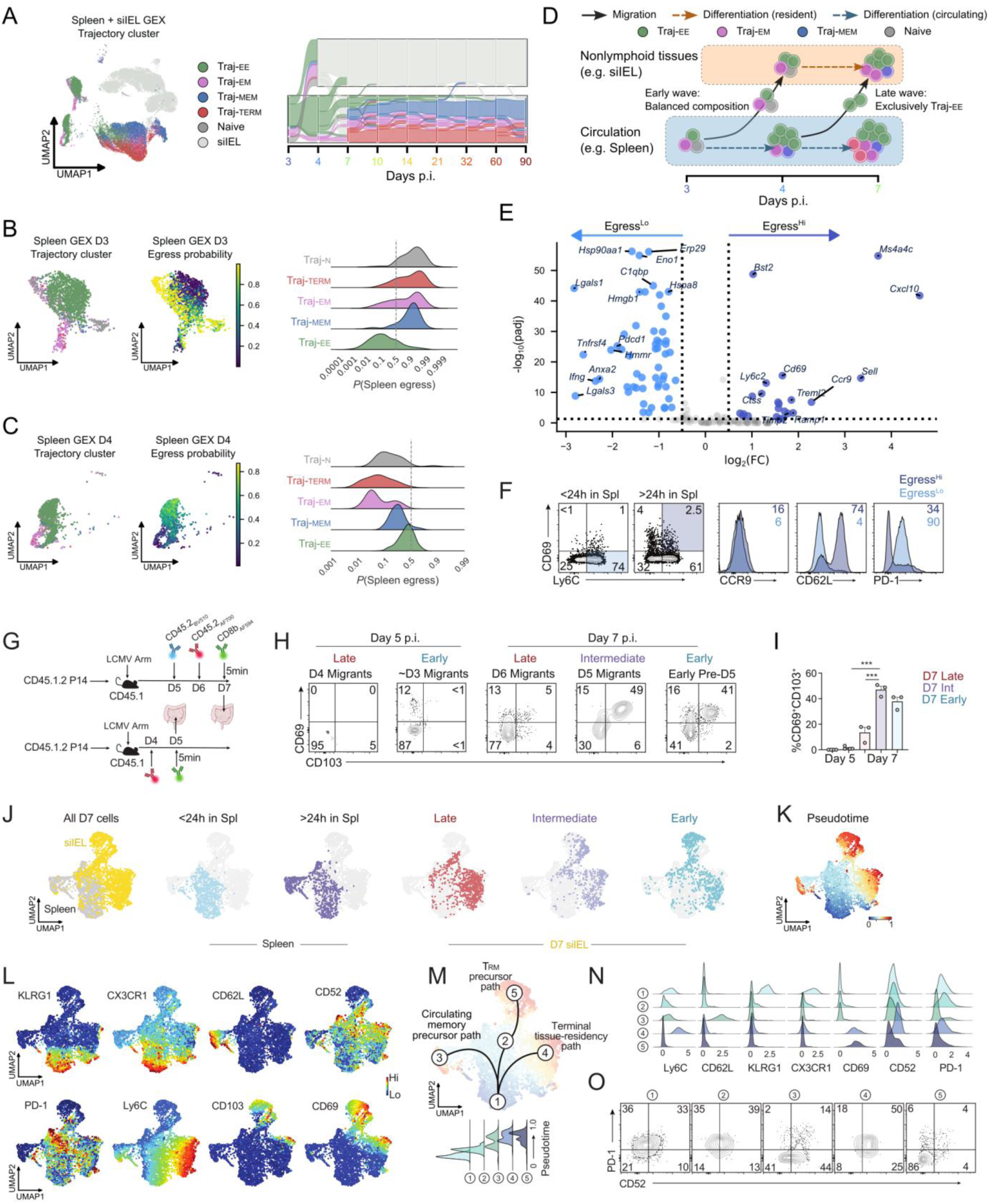
Migration kinetics influence differentiation of Trm precursors. **(A)** Circulating CD8 T cell trajectories plotted in multi-compartment UMAP space (left) and Sankey diagram (right). **(B-C)** Tissue egress probabilities for splenic P14 cells at day 3 p.i. (B) and day 4 p.i. (C). UMAPs of spleen trajectory clusters (left) and migration probabilities (middle) with histograms (right) displaying distribution of migration probability by trajectory. **(D)** Schematic summarizing OT-guided prediction of composition of cells within different waves of cells trafficking from circulation into the siIEL. **(E)** Volcano plot of differentially expressed cell surface genes in egress^hi^ cells in the spleen on day 3 p.i.. FC: fold change; padj: Benjamini-Hochberg adjusted p-values. **(F)** CD45.2_AF700_ antibody was administered to LCMV Arm infected mice on day 6 p.i. and spleens were harvested on day 7 p.i. Representative flow cytometry plots highlighting egress^lo^ and egress^hi^ P14 cell populations in the spleen on day 7 p.i. Pregating was performed based on intravasular antibody labeling and shaded gates drawn based on marker genes in E (left). Frequency of CCR9, CD62L, and PD-1 expression on egress^lo^ and egress^hi^ populations (right). **(G)** Schematic of intravenous labeling experiment to distinguish early vs. late arrivals in spleen and siIEL on day 7 p.i. (top). In a separate cohort of mice, CD45.2 antibody was administered on day 4 p.i. with spleens and siIEL harvested on day 5 p.i. **(H-I)** Representative flow cytometry plots and quantification of CD69^+^CD103^+^ siIEL cells (from G). “Early” arrival cells correspond to antibody unlabeled cells at day 7 p.i. and late arrivals represent antibody-labeled cells on day 7 p.i. **(J)** UMAPs of high-dimensional spectral flow cytometry data of day 7 p.i. cells from G. **(K)** Wishbone speudotime analysis of spectral flow cytometry data from G. **(L)** Feature plots of protein expression (bottom) shaded by fluorescence intensity on day 7 p.i. P14 cells from G. **(M)** Five clusters (from Figure S2H) are highlighted in the pseudotime-labeled UMAP space with putative differentiation paths represented by black lines (top). Speudotime scores for each cluster are plotted (below). **(N)** Expression levels of key molecules on five clusters from M. **(O)** Representative flow cytometry plots of CD52 and PD-1 expression on the following pre-gated cells corresponding do the five clusters from M: 1) <24h in spleen and Ly6c^+^CD69^-^, 2) <24h in siIEL, 3) >48h in spleen and Ly6c^+^CD69^+^CD62L^+^, 4) <24h in siIEL, and 5) >48h in siIEL and CD69^+^CD103^+^. Graphs show mean+/-SEM, n=3-4/group, student’s t test, *p<0.05, **p<0.005, ***p<0.001, (F-O).

We verified that trajectories were not systematically altered in the multi-compartment context compared with the spleen alone by means of a trajectory embedding analysis informed by spleen-only findings (Figure 2C-G). We found that dimensionality reduction on the TKME representations integrated timepoints within each compartment while still distinguishing the spleen from the siIEL, implying that long-term trajectories within each compartment were preserved and distinct in the multi-compartment model (Figure S2D). We assessed this further by clustering on the trajectory embeddings and visualizing the resulting fate flows (Figure S2E), confirming that trajectories partitioned by compartment outside of the days 3-7 p.i. migration window (Figure S2F). Finally, we assessed the degree of overlap between trajectory clusters fit on the combined model with those fit on either the spleen-only model or a siIEL-only model, finding excellent agreement between trajectories in both cases (Figure S2G). This strengthens the applicability and interpretability of our subsequent migration trajectory analysis.

The early wave of migration (days 3-4 p.i.) was composed of a relatively balanced mixture of CD8 T cell fates, whereas the later wave (days 4-7 p.i.) was dominated by Traj-ee cells (Fig. 4A). Consistent with this shift, Traj-ee cells exhibited the lowest per-cell propensities for spleen egress during the early wave (Figure 4B) but became the most egress-prone subset in the spleen during the later migration wave (Figure 4C). These observations suggest a dynamic model of CD8 T cell trafficking in which cells with diverse fates preferentially seed tissues early, while Traj-ee cells increasingly contribute to tissue entry as infection progresses (Figure 4D). This model is consistent with prior reports showing that CD127^+^ or KLRG1^-^ CD8 T cells infiltrating non-lymphoid tissues are more likely to give rise to long-lived Trm populations^55,58,59^, and extends these findings by quantitatively linking migration timing, differentiation path, and per-cell egress propensity.

To determine whether we could experimentally distinguish cells with a high propensity for egress in the spleen, we next defined gene expression of P14 cells predicted to traffic from the spleen to the siIEL in the early day 3 p.i. (termed egress^Hi^) wave relative to splenocytes less likely to migrate in this initial wave (termed egress^Lo^). We identified cells with a high probability of migration from circulation to tissue (i.e. *P*(spleen egress) > 0.5) at day 3 p.i. relative to day 4 p.i. and performed differential expression analysis (Figure 4E). Early egress^Hi^ cells exhibited elevated expression of tissue-residency–associated genes, including *Cd69* and siIEL homing chemokine receptor *Ccr9*. Egress^Lo^ cells were enriched for activation- and effector-associated transcripts such as *Pdcd1, Lgals1, Lgals3*, and *Ifng*. These transcriptional differences support a model in which early waves of tissue entry may already be enriched for cells already biased toward Trm differentiation (for example upregulating expression of CD69 prior to siIEL homing and lodging), while egress^Lo^ cells consist of terminally differentiated effector cells with reduced capacity for durable residency and memory.

To experimentally validate these predictions, we used time-resolved intravascular antibody labeling to distinguish splenic CD8 T cells that had resided in the spleen for less than or greater than 24 hours. We reasoned that cells retained in the spleen for longer periods would be more likely to acquire migration-competent programs, as interactions with antigen-presenting cells and stromal cues that instruct tissue egress are unlikely to occur immediately. Consistent with this hypothesis, we observed a higher frequency of CD69^+^Ly6C^+^ cells among splenic CD8 T cells localized to the spleen for more than 24 hours, matching the transcriptional profile of the predicted egress-high population (Figures 4E-F).

We next compared the phenotype of cells within the spleen. Egress^Hi^ cells exhibited increased expression of CCR9 and CD62L and reduced PD-1 expression, closely aligning with the OT-inferred transcriptional signatures (Figure 4F). Although Ly6C and CD62L are classically associated with lymphoid homing/localization, both markers are also linked to stem-like or less terminally differentiated states^60,61^. These observations suggest that circulating precursors of Trm may adopt a flexible, hybrid phenotype that retains stem-associated features while acquiring tissue-migration competence, potentially enabling adaptability in downstream differentiation decisions.

### Early-arriving Trm precursors undergo a distinct differentiation trajectory

We next integrated OT-guided trajectory inference with time-resolved *in vivo* labeling to further resolve early differentiation paths of Trm precursors. Using a longitudinal antibody-labeling strategy analogous to Figure 3K, we distinguished early and late tissue arrivals at day 7 p.i. (Figure 4G). In addition, we included a cohort in which circulating P14 cells were labeled at day 4 p.i., enabling identification of siIEL P14 cells at day 5 p.i. that represent some of the earliest immigrants into the intestine, with intravenously unlabeled cells presumed to have arrived between days 3-4 p.i.

Time-resolved labeling revealed a progressive tissue-residency differentiation program marked by stepwise upregulation of CD69 and CD103 in the siIEL (Figure 4H-I). Leveraging the robustness of this labeling approach, we next focused on distinct waves of migrating cells in both the spleen and intestine at day 7 p.i. UMAP visualization of spectral flow cytometry data showed clear segregation of splenic and siIEL populations, as expected, with additional clustering driven by tissue localization and migration timing (Figure 4J). Notably, Wishbone pseudotime analysis closely aligned with time-resolved labeling and recapitulated the differentiation paths predicted by OT-based modeling (Figures 3K–N and 4E), supporting a coherent temporal progression toward tissue residency (Figure 4K).

We next examined phenotypic changes in P14 cells as they progressed along these early differentiation trajectories by integrating surface marker expression, unbiased clustering, and pseudotime analysis (Figure 4K-N, S2H). Recent migrants in the spleen exhibited elevated KLRG1 and CX3CR1 expression, consistent with recently primed effector cells representing the earliest post-activation stage in this model (Figure 4L-M). In contrast, P14 cells progressing toward a CD69^+^CD103^+^ Trm fate in the siIEL progressively downregulated CX3CR1 and KLRG1 and transiently upregulated CD52 and PD-1 (cluster #2, Figure 4L-N). As CD69 and CD103 expression increased in these cells, CD52 and PD-1 levels declined, consistent with stabilization of a mature tissue-resident program (cluster #5, Trm precursors).

We also identified an alternative differentiation route within the siIEL characterized by sustained CD52 and Ly6C expression (Figure 4M–N). Cells that failed to downregulate Ly6C were biased toward a short-lived tissue-resident fate or potential re-entry into circulation (cluster #4). These transitions are summarized in a putative differentiation model (Figure 4M) and visualized as expression histograms (Figure 4N). Guided by pseudotime and clustering analyses, we further validated these populations using hierarchical flow cytometry gating, confirming distinct CD52 and PD-1 expression patterns across trajectories (Figure 4O). Together, these data define early bifurcation points following tissue entry and reveal a temporally ordered phenotypic progression that distinguishes long-lived Trm precursors from short-lived or transient tissue migrants.

### Time of tissue entry stratifies long-lived Trm subsets along a CD52-CD69-CD103 axis

Having established that early-arriving CD8 T cells entering the siIEL are biased toward long-lived tissue-resident fates, we next examined how differentiation unfolds after tissue entry and how long-lived Trm heterogeneity emerges over time. We fit an OT trajectory model restricted to the siIEL compartment, accounting for proliferation and apoptosis in the source marginals as in our prior analyses (Figures 5A, S3A). Consistent with experimental observations^12^, inferred siIEL population kinetics peaked at day 7 p.i. and declined thereafter (Figure S3B). Quality-control analyses confirmed that OT inference was not confounded by technical artifacts or cell cycle effects (Figure S3C-D).

**Figure 5.**
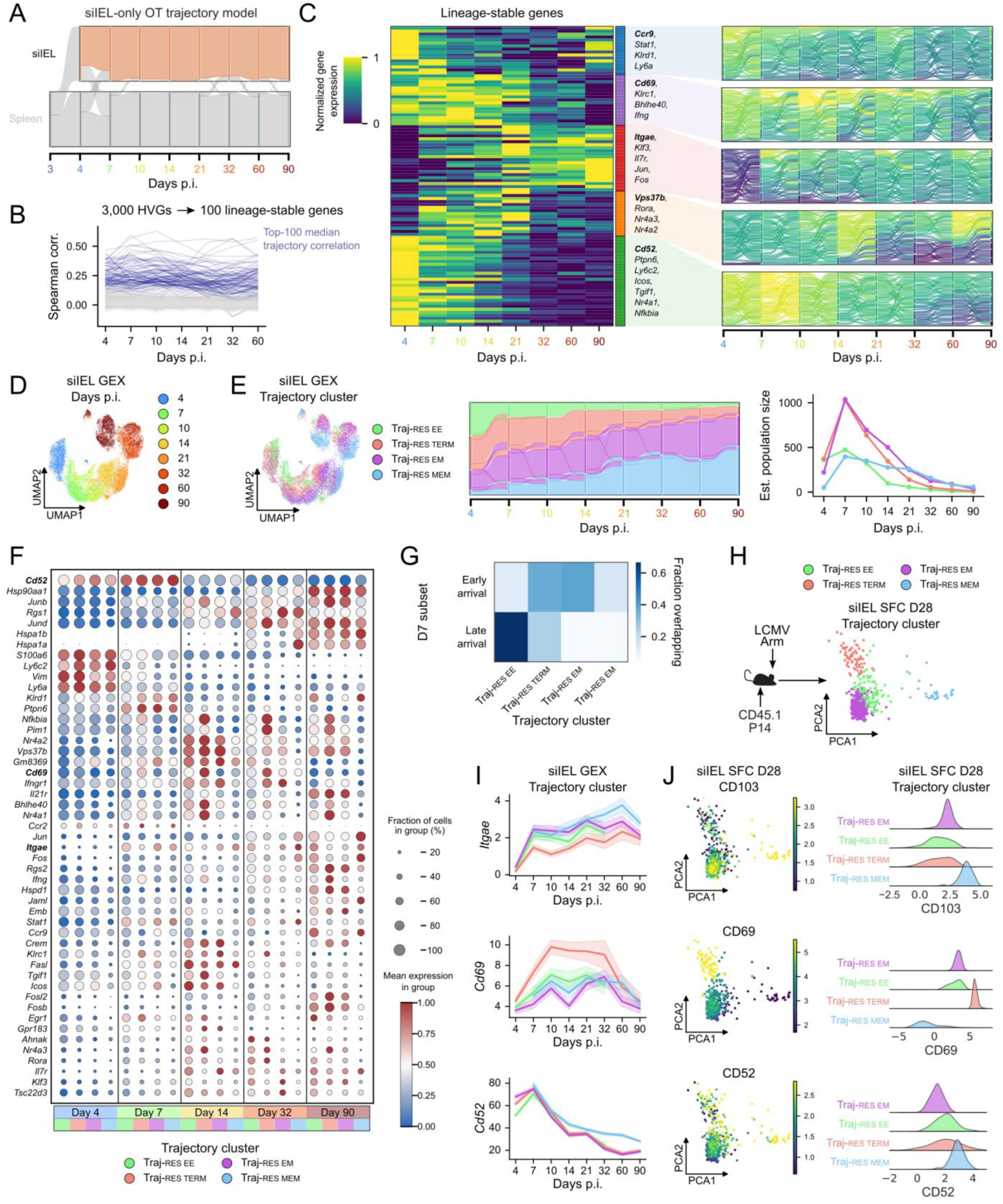
Time of tissue entry stratifies long-lived Trm subsets along a CD52-CD69-CD103 axis. **(A)** Multi-compartment OT-modeling with only siIEL resident cells highlighted **(B)** Traces of gene expression correlations over time in siIEL cells, induced by OT mappings. The top 100 genes with highest median correlations are shaded blue. **(C)** Heatmap of normalized mean gene expression for the top 100 lineage-stable genes (left) and select Sankey diagrams (right) colored by supercell expression for representative lineage-stable genes from each module. **(D)** UMAP of siIEL CD8 T cells shaded by timepoint. **(E)** siIEL-specific trajectory clusters projected in GEX UMAP space (left), Sankey diagram (middle), and population kinetics reconstruction (right). **(F)** Dotplot of gene expression for trajectory clusters at representative time points (days 4, 7, 14, 32, and 90 p.i.). Genes correspond to top lineage genes derived from Figure 5B, filtered to display genes encoding cell-surface proteins, secreted proteins, or transcription factors. **(G)** Confusion matrix displaying overlap between D7 siIEL migrant populations based on arrival time (rows) and trajectory clusters (columns), normalized by row. **(H)** Congenic P14 cells were transferred to mice subsequently infected with LCMV Arm. On day 28 p.i., high-dimensional spectral flow cytometry analysis was performed on siIEL cells and PCA was performed. OT-guided trajectories were mapped to P14 cells based on gene expression patterns of CD69, CD103, and CD52. **(I)** Line plots of mean gene expression for *Itgae, Cd69*, and *Cd52* over time, grouped by trajectory cluster (from E). Shaded regions represent 95% confidence intervals. **(J)** PCA was performed on spectral flow data from H and shaded by degree of marker expression of CD103, CD69, and CD52 with corresponding histograms right.

To identify gene programs that structure Trm differentiation over time, we first selected the 100 genes with the strongest trajectory correlations and clustered them by temporal expression dynamics (Figure 5B-C). Canonical Trm markers exhibited distinct but temporally broad patterns: *Cd69* peaked early (around day 10 p.i.), whereas *Itgae* (CD103) increased gradually through late memory timepoints (note: this is relative to only siIEL cells, both *Cd69* and *Itgae* are near-uniformly higher compared to splenocytes). Although these genes clearly reflect tissue-residency identity, they did not by themselves resolve stable differentiation paths. Similarly, stemness and terminal differentiation scores varied across time but weakly distinguished Trm subsets (Figure S3E). In contrast, genes such as *Cd52* and *Vps37b* displayed strong trajectory correlations and segregated fate flows into distinct differentiation paths, suggesting previously underappreciated aspects of Trm heterogeneity (Figure 5C).

To better resolve Trm differentiation paths, we performed trajectory-based embedding and clustering of siIEL P14 cells. Whereas GEX-based embeddings were dominated by timepoint (Figure 5D), TKME-integrated cells revealed four stable trajectory clusters with high internal consistency (Figures 5E, S3F–G). Based on population persistence and GEX profiles, we labeled these trajectories as resident early effector (Traj-res ee), resident terminal effector (Traj-res-term), resident effector memory (Traj-res-em), and resident memory (Traj-res-mem), ordered by increasing longevity.

Population dynamics inferred from OT predicted that Traj-res-ee and Traj-res-term dominate the siIEL during the acute phase (days 4-7 p.i.), coinciding with peak viral burden, whereas Traj-res-em and Traj-res-mem progressively accumulate and persist into memory timepoints (Figure 5E, right). These identities were supported by trajectory-correlated gene expression: Traj-res-ee and Traj-res-term showed elevated *Ifng* and *Bhlhe40*, Traj-res-mem was enriched for *Jun/Fos* with low effector gene expression, and Traj-res-em exhibited intermediate profiles consistent with a transitional fate (Figure 5F).

We next related these trajectories to migration timing and prior Trm classification schemes^13,28^. Mapping day 7 p.i. early and late arrivals (Figure 3I) onto siIEL trajectories revealed that late-arriving cells preferentially populated Traj-res-ee and Traj-res-term, whereas early arrivals seeded the long-lived Traj-res-em and Traj-res-mem trajectories, along with a minority of Traj-res-term cells (Figure 5G). This pattern mirrors our earlier findings that early tissue entry favors stem-like and persistent fates, whereas later entry is biased toward transient effector programs. Consistent with this interpretation, Traj-res-mem cells exhibited higher gene set scores for previously defined stem-like Trm programs^13,28^ and lower scores for terminal Trm programs^13,28^ at late timepoints, whereas the other trajectories showed the opposite pattern (Figure S3H). Thus, trajectory-based clustering not only recapitulates established Trm classifications but also clarifies their developmental relationships and timing of specification.

Finally, we sought experimentally tractable markers that distinguish Trm trajectories *in vivo*. Traj-res-mem cells upregulated both *Cd52* and *Itgae*, whereas Traj-res-term cells showed high *Cd69* and low *Itgae*, with Traj-res-ee and Traj-res-em occupying intermediate states (Figure 5F). These expression patterns were stable over time (Figure 5F). High-dimensional spectral flow cytometry of siIEL P14 cells at day 28 p.i. revealed discrete populations defined by CD103, CD69, and CD52 expression (Figure 5H-J), enabling direct identification of these trajectories at the protein level. Notably, co-expression of CD52 and CD103 robustly marked the long-lived Traj-res-mem population of cells, providing a practical strategy for investigating stem-like Trm cells *in vivo*.

### Context-aware trajectory analysis reveals shared and compartment-specific programs governing circulating and resident CD8 T cell differentiation

OT-based fate mapping provides a natural framework for comparing differentiation programs across anatomical compartments, where CD8 T cell states are related but not identical. Although circulating and tissue-resident populations are transcriptionally and phenotypically distinct, both generate long-lived memory cells. We therefore used OT-derived trajectories to distinguish shared versus tissue-specific differentiation programs, focusing our comparisons on fate rather than static state (Figure 6A).

**Figure 6:**
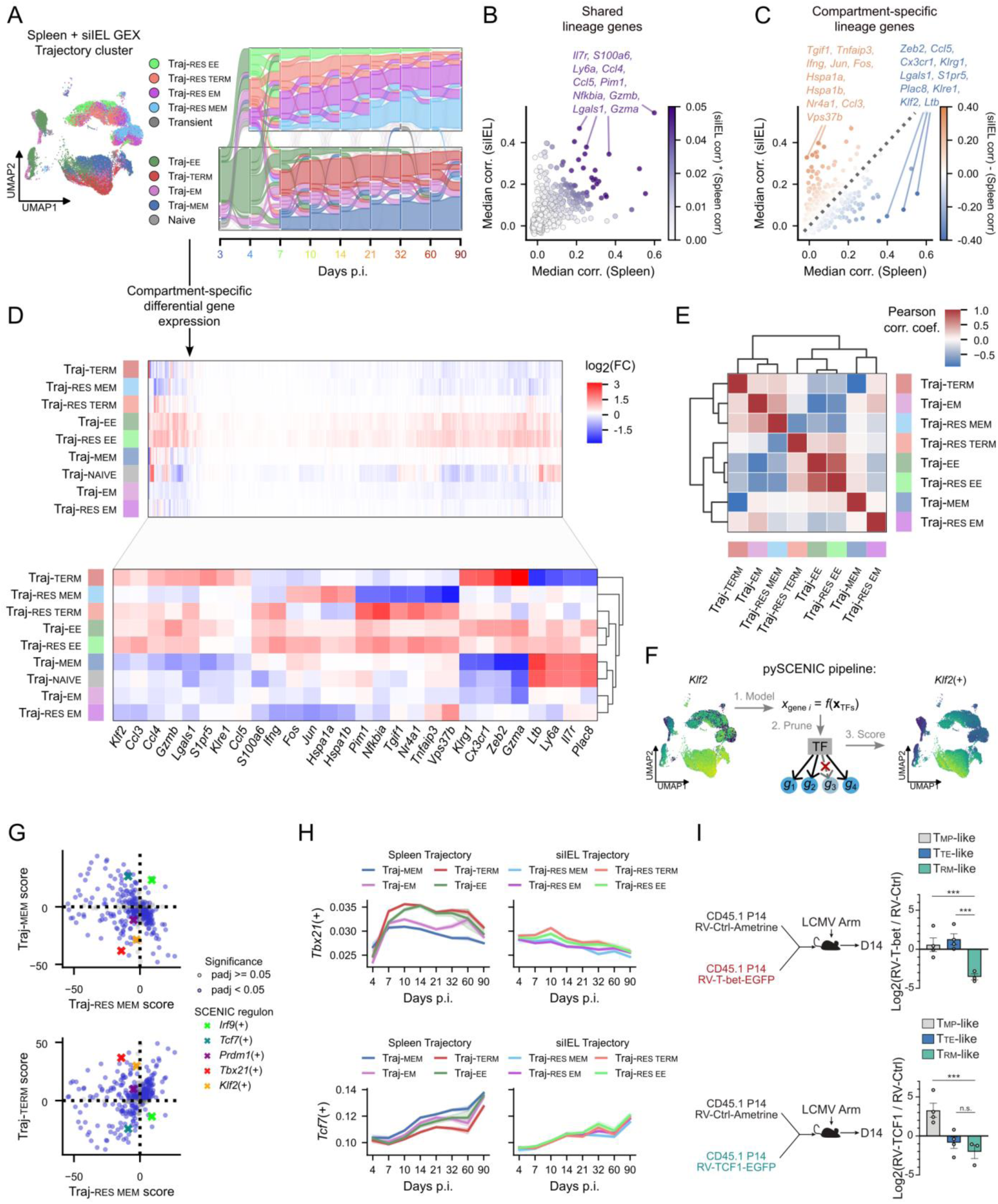
Context-aware trajectory analysis identifies shared and distinct differentiation programs in circulating and resident trajectories. **(A)** Schematic of comparative analysis approach integrating both circulating and tissue-resident OT-informed trajectories. **(B-C)** Scatterplots of median trajectory correlation (spearman) for genes in the siIEL vs. spleen trajectories, colored by product of correlations (B) and difference between siIEL and spleen correlations (C). Top shared genes (B) and siIEL/spleen-specific genes (C) genes are highlighted. **(D)** Heatmaps of log_2_-fold changes for spleen and siIEL trajectory clusters, with all highly variable genes (top) and top shared and compartment-specific genes highlighted (bottom). **(E)** Heatmap showing Pearson correlation coefficients between trajectory cluster log_2_-fold changes within spleen and siIEL compartments. **(F)** Schematic of SCENIC method for fitting transcription factor regulons. *Klf2* regulon prediction is shown as an example. **(G)** Scatterplots of Traj-mem and Traj-term vs. Traj-res-mem with t-statistics for SCENIC regulons. Regulons for *Irf9*(+), *Tcf7*(+) *Prdm1*(+), *Tbx21*(+), and *Klf2*(+) are highlighted. **(H)** Regulon dynamics of *Tbx21*(+) and *Tcf7*(+) are plotted. **(I)** Congenic P14 cells transduced with control retrovirus (RV-Ctrl), T-bet-encoding retrovirus (RV-T-bet), or TCF1-encoding retrovirus (RV-TCF1) were transferred into mice infected with LCMV Arm. Spectral flow cytometry quantification of canonical cell states and phenotypes are reflected. Graphs show mean+/-SEM, each dot represents an individual mouse, students t test, n=3-5/group, ***p<0.005, n.s. non-significant.

From the top 3,000 highly variable genes identified in spleen and siIEL datasets, we retained 1,816 genes shared between compartments. For each gene, we compared its trajectory correlation under spleen- and siIEL-specific OT models, reflecting how stably gene expression is propagated along inferred fate paths (Figures 1M and 5B). This analysis revealed both shared lineage genes and compartment-specific programs (Figures 6B-C). Shared genes included canonical regulators such as *Il7r, Ly6a, Ccl4, Ccl5, Pim1, Nfkbia, Gzmb*, and *Gzma* (Figure 6B), indicating conserved roles in CD8 T cell differentiation. In contrast, spleen-enriched genes (*Zeb2, Cx3cr1, Klrg1, S1pr5, Klf2*) and siIEL-enriched genes (*Tgif1, Tnfaip3, Ifng, Jun, Nr4a1, Vps37b*) highlighted compartment-specific differentiation programs (Figure 6C).

We next examined how these programs were deployed across trajectory clusters. Differential expression analysis controlling for timepoint revealed that early effector trajectories in both compartments (Traj-ee and Traj-res-ee) exhibited globally elevated gene expression, consistent with acute activation of these cells (Figure 6D). Memory trajectories showed both overlap and divergence: *Jun* and *Fos* were upregulated in Traj-mem and Traj-res-mem, while *Il7r* was shared between Traj-mem and Traj-res-em, suggesting partial reuse of memory-associated programs. To quantify cross-compartment similarity, we correlated trajectory-specific log-fold changes (Figure 6E). Early effector trajectories were highly conserved between compartments, whereas memory trajectories diverged substantially between circulating and resident paths. Circulating memory (Traj-mem) showed weak positive correlations with multiple siIEL trajectories, while Traj-em bifurcated toward terminal-like versus resident-memory-like programs. These patterns suggest that circulating memory differentiation follows a broadly permissive, stem-like program, whereas tissue-resident cells engage a distinct transcriptional axis more closely aligned with effector-memory states.

To define the regulatory architecture underlying these differences, we applied SCENIC^62,63^ to infer transcription factor-centered regulons (defined as transcription factors and the set of genes that they regulate) and tested their association with trajectories while controlling for timepoint (Figure 6F). Effector-associated regulons were largely conserved across compartments. In particular, the *Tbx21*(+) (positive regulation by T-bet) regulon was reduced across both circulating and resident memory trajectories, while it was elevated in Traj-term cells (Figure 6G). *Tbx21*(+) activity peaked around day 10 p.i. in effector trajectories and declined thereafter, and declined more rapidly in the siIEL than in the spleen while remaining low in memory trajectories (Figure 6H, top). Accordingly, enforced T-bet expression induced accumulation of Tte-like cells relative to Trm cells at day 14 p.i. (Figure 6I, top), establishing T-bet as a repressor of Trm differentiation in the gut, in line with reported roles in the skin and lung^64–66^.

In contrast, memory-associated regulons displayed strong context dependence. The *Tcf7*(+) regulon was prominent in circulating memory but markedly attenuated in tissue-resident trajectories (Figure 6G). Memory regulon activity diverged substantially among spleen trajectories but was compressed in the siIEL (Figure 6H, bottom), suggesting that tissue-residency constrains canonical memory programs. Consistent with this, TCF1 overexpression increased frequencies of Tmp-like cells at day 14 p.i., while decreasing the frequency of Trm in the siIEL (Figure 6I, bottom). These observations support a model in which tissue-residency alters canonical memory programs (e.g. driven by TCF1), constraining their utilization during Trm differentiation.

Given that both T-bet and TCF1 expression restrained Trm formation and exhibit dynamic regulons in spleen and siIEL trajectories, we next investigated their role in tuning early Trm fate decisions (Figure S4A). In the siIEL, we found that sustained T-bet expression promoted an effector-like CX3CR1^hi^PD-1^lo^ phenotype (Figure S4B), corresponding to the initiation of the pseudotime differentiation trajectory (Figures 4M, S4C). Interestingly, forced T-bet expression also promoted a memory-like CX3CR1^lo^CD62L^hi^ state in the spleen (Figure S4B), suggesting that T-bet may play dual roles inhibiting differentiation towards Trm while promoting circulating memory precursors early in infection (Figure S4C). TCF1 overexpression promoted a more mature CX3CR1^lo^PD-1^lo^ phenotype in the siIEL as well as a CX3CR1^lo^CD62L^hi^ phenotype in the spleen (Figure S4B), situating TCF1 as an early potentiator of both circulating and Trm precursor paths (Figure S4C). This early timepoint illustrates temporal dependence of transcription factor regulation and motivates further study into context-dependent gene regulation during different windows of the infection response.

These results suggest effector differentiation is governed by similar regulatory programs across tissues, whereas memory differentiation is more strongly shaped by local tissue context. Circulating memory cells appear to maintain canonical stem-like programs, while tissue-resident memory cells adopt a distinct regulatory state influenced by the tissue environment. By modeling differentiation as dynamic fate flows across compartments, the OT framework enables systematic identification of both shared and tissue-specific mechanisms that shape CD8 T cell fate.

### Tissue-specific regulatory architecture of effector and memory specification

Finally, we used our combined profiling of circulating and tissue-resident CD8 T cell trajectories to examine the broader regulatory landscape of the CD8 T cell response during acute infection. Many previous studies have compared circulating and tissue-resident CD8 T cells, often relying on static endpoint phenotypes or intermediate transcriptional states. While these approaches have been instrumental in defining Trm identity, they provide less insight into how regulatory programs unfold over time or diverge along distinct differentiation paths. By pairing longitudinal sampling with trajectory - informed and regulon-based analyses, our approach allows us to follow how TF programs are engaged across compartments and subsets.

Rather than focusing on individual differences between circulating and resident cells, we aimed to organize regulons into broader modules based on their compartment- and cluster-specific activity, with the goal of identifying regulatory programs that operate across the CD8 T cell differentiation landscape. We filtered SCENIC regulons for robust trajectory associations and computed mean regulon activity (AUCell) across timepoints and subsets. To minimize confounding from ongoing tissue migration, we restricted this analysis to timepoints from day 7 p.i. and beyond. Unsupervised clustering of regulons by their standardized activity profiles uncovered distinct temporal and subset-specific modules of TF activity (Figure 7A). To contextualize the functional relevance of these modules, we examined regulon activity along three axes: overall activity in spleen versus siIEL (Figure 7B), cluster-specific activity within the spleen (Figure 7C), and cluster-specific activity within the siIEL (Figure 7D). This multi-dimensional framework enabled us to distinguish regulons that are broadly active across compartments from those whose function is more context dependent. Representative regulons from each module revealed clear compartmental biases, with *Tcf7*(+), *Tbx21*(+), and *Tfap4*(+) (AP4) modules consistently exhibiting higher activity in the spleen, while the *Bcl6*(+) module was preferentially active in the siIEL (Figure 7B). The *Foxp1*(+) module displayed intermediate behavior, with elevated activity in the siIEL through day 90 p.i. followed by a gradual increase in the spleen, suggesting delayed engagement of this regulatory program in circulation.

**Figure 7:**
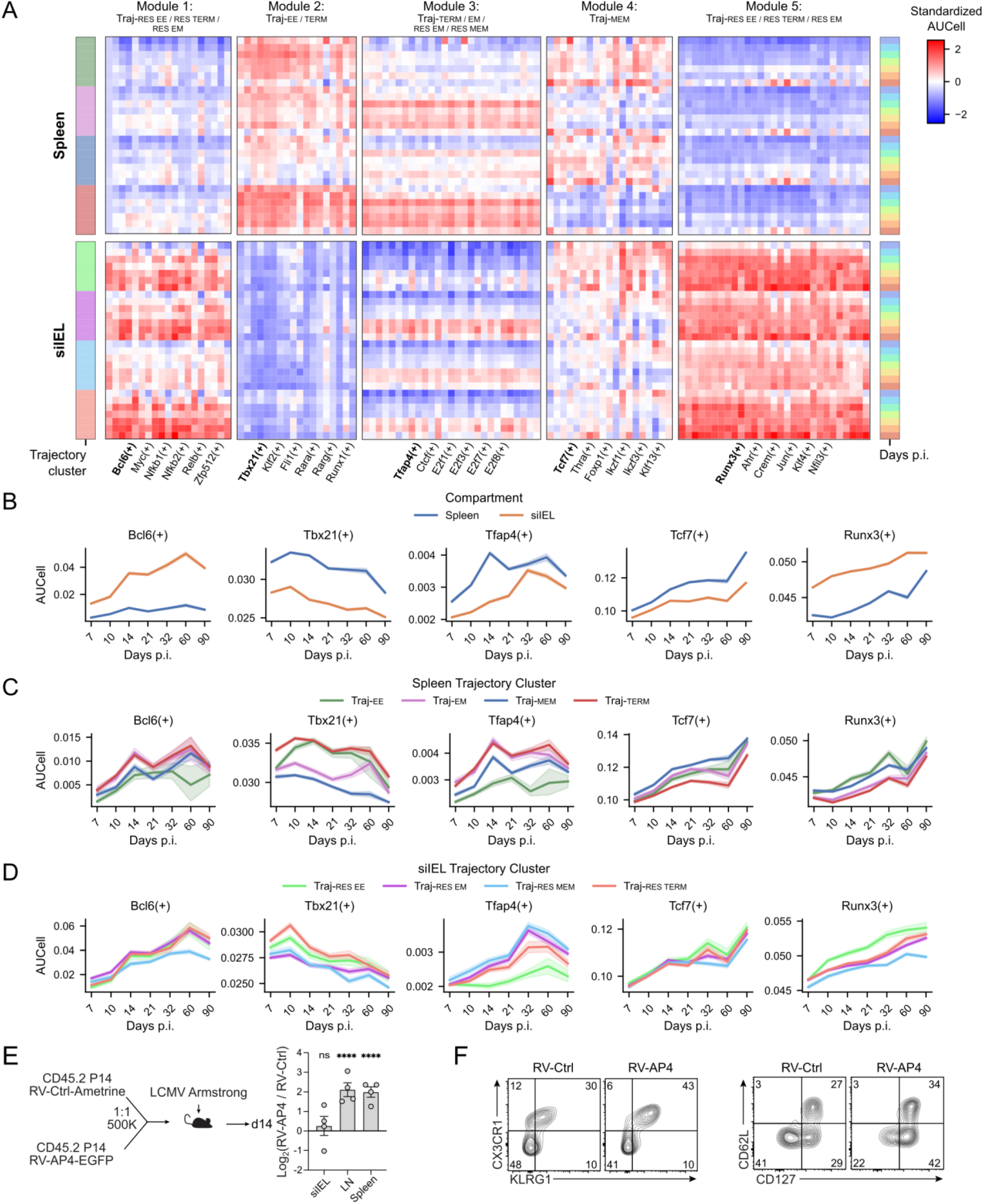
Compartment-specific regulatory architecture of effector and memory specification. **(A)** Heatmap of standardized regulon activity (AUCell), grouped by compartment, time point, and trajectory. **(B-D)** Mean AUCell score over time for representative regulons from each module, with hue corresponding to average AUCell score in spleen vs. siIEL (B), spleen trajectory clusters (C), and siIEL trajectory clusters (D). **(E)** Congenic P14 cells transduced with control retrovirus (Ctrl) or AP4-encoding retrovirus were mixed 1:1 and transferred into mice infected with LCMV Arm. Donor P14 cells were enumerated in the spleen, mesenteric lymph node (LN), and siIEL on day 14 p.i. ( **F)** Representative spectral flow cytometry plots for CX3CR1, KLRG1, CD127, and CD62L for donor splenocytes are shown. Graphs show mean+/-SEM, n=4/group, student’s t test, ****p<0.0005 relative to siIEL.

Despite these compartmental differences, several modules exhibited conserved temporal dynamics. *Tcf7*(+) activity increased over time, whereas *Tbx21*(+) activity declined, consistent with their established roles in memory versus effector differentiation. However, while *Tbx21*(+) activity was consistently enriched in effector subsets across both compartments, *Tcf7*(+) showed greater heterogeneity among spleen clusters than in the siIEL, reinforcing the earlier conclusion that memory differentiation may be more context restricted in tissue-resident cells.

In contrast, the *Tfap4*(+) module displayed compartment-specific kinetics. *Tfap4*(+) activity rose rapidly in the spleen between days 7 and 14 p.i. before stabilizing, whereas in the siIEL it increased more gradually between days 10 and 32 p.i. before converging on similar activity levels as the spleen (Figure 7B-D). Cluster-level analysis predicted that *Tfap4*(+) activity was highest in Traj-term and Traj-em paths in the spleen, as well as in Traj-res-em and Traj-res-mem paths in the siIEL (Figure 7C–D). This pattern suggested partial overlap between the regulatory programs controlling circulating effector-memory and tissue-resident memory differentiation. AP4 deficiency has previously been shown to limit proliferation and metabolism of circulating effector CD8 T cells^67^. However, the role of AP4 in governing tissue-resident cells remains unclear and whether augmenting AP4 expression levels modulates memory formation in infection has not been investigated. To test this prediction experimentally, we overexpressed AP4 in P14 cells in a mixed transfer setting in LCMV Arm (Figure 7E, left). AP4 overexpression enhanced the accumulation of early memory populations in circulation in the spleen and mesenteric lymph node, but failed to alter the abundance of tissue-resident cells in the intestine (Figure 7E, right), consistent with its predicted regulon patterns at earlier infection timepoints (Figure 7B). Further, AP4 overexpression promoted effector-associated phenotypes in circulating CD8 T cells, including increased frequencies of KLRG1^+^CX3CR1^+^ Tte-like cells and CD127^+^CD62L^−^ Tem-like cells in the spleen (Figure 7F). Together, these results identify AP4 as a context-specific regulator that preferentially amplifies circulating effector CD8 T cell responses.

## DISCUSSION

In this study, we developed an OT-based framework to reconstruct CD8 T cell fate flows across time and tissue during viral infection. By explicitly accounting for population expansion, contraction, and migration, our approach addresses several practical limitations of trajectory inference in highly non-equilibrium immune responses. Integrating circulating and tissue-resident compartments, we provide a unified “fate-flow” view of CD8 T cell differentiation and highlight early, context-dependent events that ultimately shape subsequent effector and memory outcomes.

We first applied OT to circulating CD8 T cells, where the underlying biology has been studied extensively. Incorporating proliferation and apoptosis signatures allowed us to recover the expected expansion-contraction kinetics of the antiviral response, supporting the idea that OT captures population-level dynamics in this setting. Beyond kinetics, OT-based fate mapping identified lineage-stable markers and resolved differentiation trajectories corresponding to canonical CD8 T cell states. Trajectory-centric clustering also helped clarify the status of effector memory (Tem) cells, which emerged as durable intermediates between terminal effector and stem-like memory programs rather than a fully separate endpoint. This provides a natural explanation for why Tem phenotypes often appear heterogeneous across studies^50^.

A key extension of this work is the move to a multi-compartment model that jointly represents differentiation and migration. Trm development has been difficult to resolve because tissue entry and Trm differentiation are asynchronous, and snapshot measurements of mixed cells are comprised of different stages of tissue adaptation. By modeling transport between spleen and siIEL, we find that time of tissue entry is strongly associated with Trm fate. Through longitudinal intravascular labeling of T cells, we show that early-arriving cells preferentially seed long-lived resident memory populations, whereas later-arriving cells display more terminal effector-like features and reduced persistence. Intravenous labeling experiments support these predictions and, importantly, show that arrival timing can imprint durable phenotypic signatures in tissue-resident CD8 T cells. These distinct stages of differentiation may be influenced by the abundance of available antigen and the complex changes in the environmental milieu influencing CD8 T cell specification. In summary, our novel application of OT to model population flow between distinct compartments provides a framework to study the interrelated roles of migration and differentiation in other biological systems.

We also used comparative analyses across compartments to connect fate flows to gene regulatory programs. Effector-associated programs were broadly conserved between circulating and resident populations, whereas memory-associated programs were more context-dependent. In our data, Trm subsets were transcriptionally closer to circulating effector memory than to central memory, consistent with emerging views that resident memory follows a distinct developmental logic. The proposed fate flow logic revealed unreported roles for both canonical and understudied TFs in regulating cell fate. Genetic perturbation experiments supported nuanced, context-specific roles for Klf2, T-bet, and TCF1 in specifying circulating and tissue-resident differentiation. Further, we show that sustained AP4 expression augments circulating memory responses but not Trm.

These results highlight the complementary utility in considering CD8 T cell differentiation in terms of a fate flow model in addition to their more widely used static labels. OT offers a practical way to infer these flows directly from independent and discrete timepoints while accommodating proliferation, death, and migration. In summary, our work establishes OT-based fate mapping as a useful framework for reconstructing immune cell dynamics and highlights migration timing as an important determinant of tissue-resident CD8 T cell specification. By bringing differentiation and migration into a single integrated model, this study provides a clearer picture of Trm formation. Beyond acute infection, CD8 T cells and other lymphocytes exhibit similar differentiation flows in chronic infection, autoimmune disease, and cancer. Fate flow modeling through robust tools such as OT may reveal key regulators of immune cell differentiation that can be harnessed to augment immunotherapies.

### Limitations of the study

We acknowledge several experimental and computational limitations. Discrete CD8 T cell states remain essential for defining phenotypes, molecular drivers, and enabling comparisons across models and infections. Rather than proposing new labels, our goal is to show how viewing T cells along continuous differentiation trajectories can reveal relationships that are obscured by static compartmentalization. At the same time, experimental perturbations necessarily act on discrete states, limiting our ability to directly manipulate entire trajectories (for example, when assessing regulators such as AP4). Furthermore, while computational OT provides probabilistic estimates for cell differentiation, our fate flow framework may misclassify ground truth lineage relationships. The *in vivo* antibody labeling strategy used here provides strong temporal resolution of migration dynamics, although very transient tissue-residency or rapid inter-tissue trafficking may not be as readily detected. Future approaches incorporating more temporally refined or iterative labeling strategies may further enhance resolution of these dynamics. Finally, our analyses predominantly focus on early fate decisions in the spleen and intestine using LCMV Arm as a model system; differentiation dynamics may vary across infections and tissue contexts. Taken together, this work combines computational inference with reductionist experimentation to introduce a conceptual framework for early CD8 T cell fate specification that we anticipate will be broadly applicable to immune cells undergoing dynamic, multi-tissue responses.

## RESCOURCE AVAILABILITY

### Lead Contact

Information and requests should be directed to the lead contacts, Justin Milner and Natalie Stanley. All materials and resources will be made available upon request.

### Material availability

All materials including used in this manuscript, including mouse lines and reagents, will be made available upon request to the lead contact.

### Data and code availability

All data are available upon request, and all original code is open source. Code for generating all figures may be found at https://github.com/alecplotkin/Optimal-Transport-CD8-Fate-Mapping. In addition, we have packaged tools for **<u>s</u>**ingle-**<u>c</u>**ell **<u>o</u>**ptimal **<u>t</u>**ransport **<u>t</u>**rajector**<u>y</u>** analysis (scotty-tools), including fate flow visualization and trajectory clustering, at https://github.com/alecplotkin/scotty-tools. No sequencing datasets were generated for this study, but publicly available datasets utilized are indicated in the key resources table.

## ACKNOWLEDGMENTS

This study was supported by NIH R00CA234430 (JJM), NIH R01A1177864 (JJM), V Foundation Scholar Award (JJM), Mary Kay Ash Foundation Award (JJM), Lung Cancer Initiative Career Development Award (JJM), Hirshberg Foundation Seed Grant (JJM), UNC Pancreatic Cancer SPORE Career Enrichment Award (JJM), American Association for Immunologists Intersect Award (WDG), UNC CGIBD Pilot Award (JJM; supported by P30 DK034987), UNC Computational Medicine Pilot Program Award (NS & JJM), University Cancer Research Fund (JJM), NIH R21AI171745 (NS), R21AG084251 (NS), UNC Bioinformatics and Computational Biology T32 Training Grant (ALP, supported by NIH 5T32GM135123-02), NIH T32-CA196589 (WDG). The UNC Advanced Analytics Core is supported by NIH P30-DK034987. The PSC is supported in part by an NCI P30CA016086.

## AUTHOR CONTRIBUTIONS

Conceptualization: A.L.P., N.S., and J.J.M.; data curation: A.L.P. and J.J.M.; formal analysis: A.L.P., G.N.M., N.S., and J.J.M.; funding acquisition: J.J.M. and N.S.; investigation: A.L.P., G.N. M., W.D.G., H.S., N.S., and J.J.M.; methodology: A.L.P., G.N.M., N.S., and J.J. M.; resources: N.S., and J.J.M.; supervision: N.S., and J.J.M.; visualization: A.L.P., G. N.M., N.S., and J.J.M.; project administration: N.S. and J.J.M.; writing – original draft: A.L.P, G.N.M., N.S., and J. J.M.; writing – review & editing: A.L.P, G.N.M., N.S., and J. J.M.

### Declaration of generative AI and AI-assisted technologies in the manuscript preparation process

During the preparation of this work the author(s) used Perplexity, powered by GPT-5.1 (Perplexity AI), in order to recommend related literature, generate and debug code for plotting figures, and parsimoniously edit text for clarity. After using this tool/service, the author(s) reviewed and edited the content as needed and take(s) full responsibility for the content of the published article.

## STAR Methods

### Key resources table

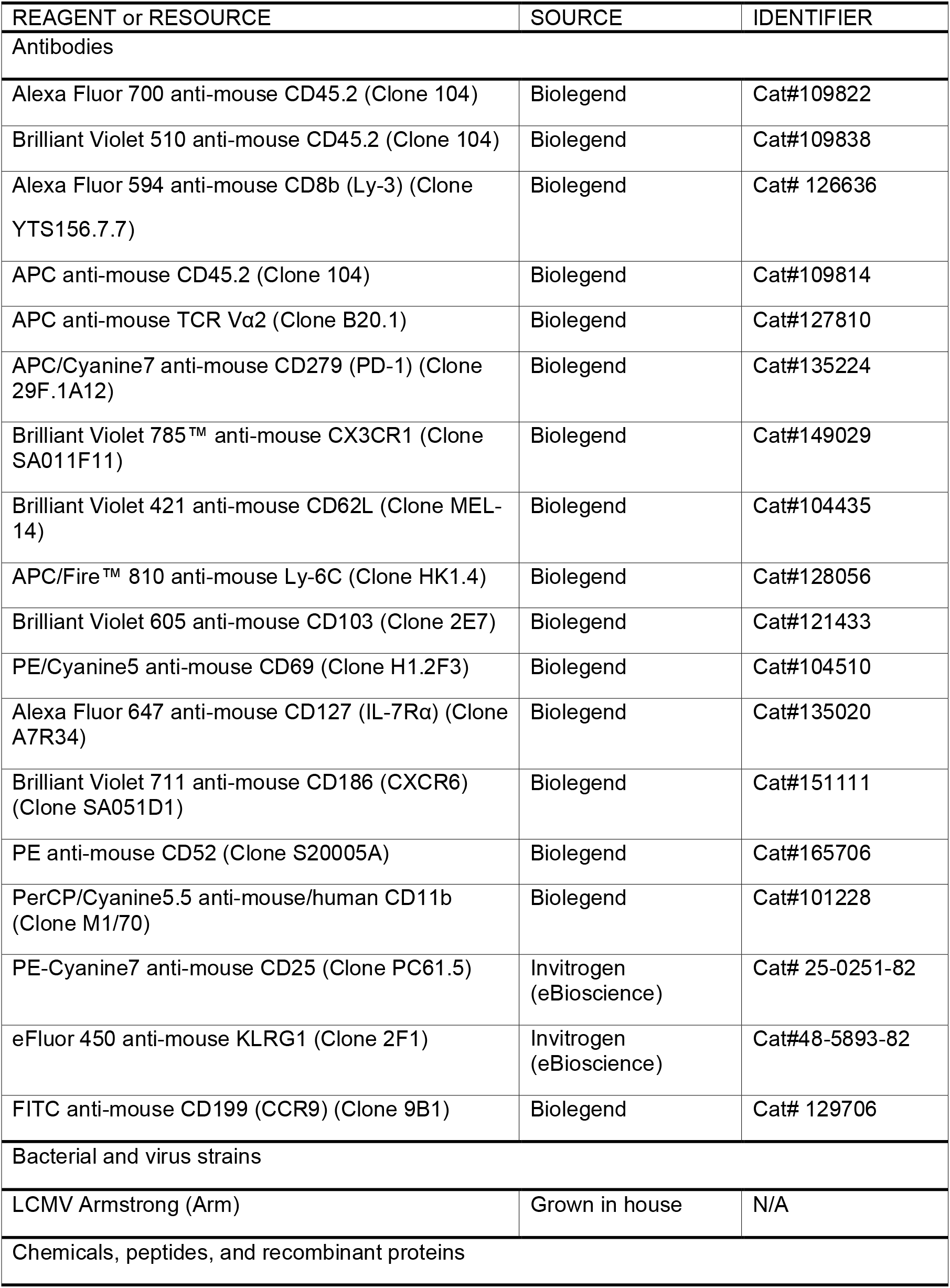
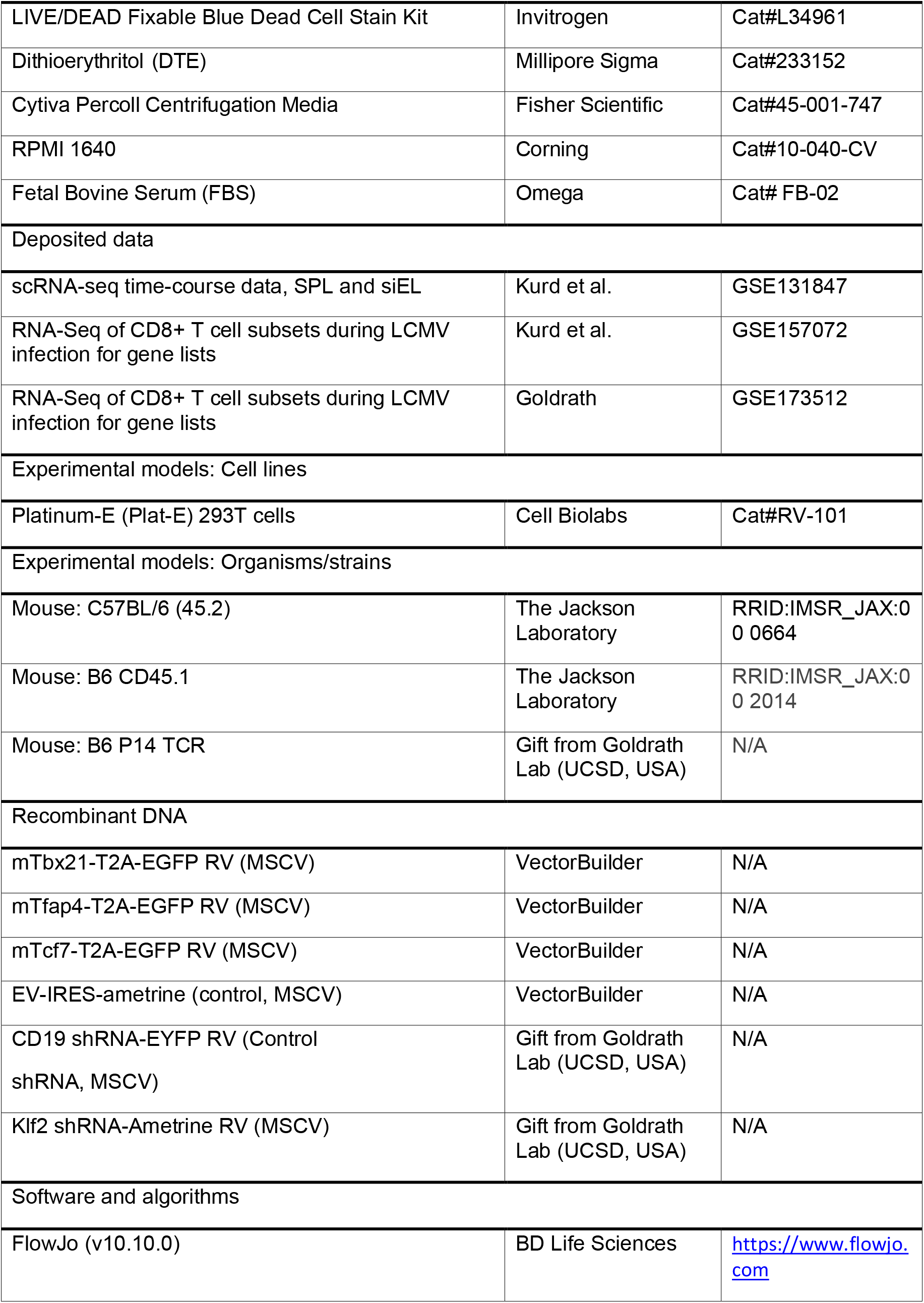
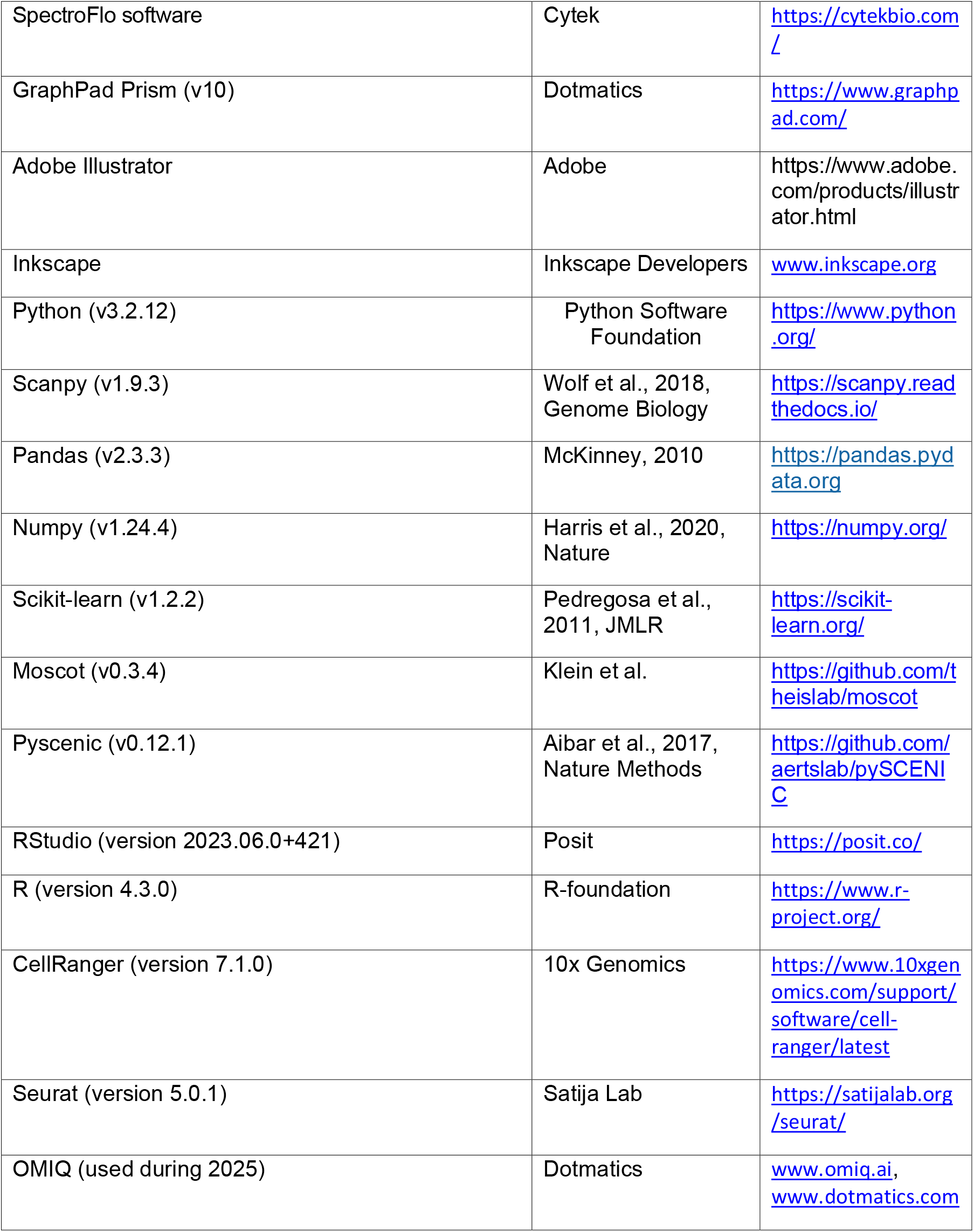

## EXPERIMENTAL MODEL DETAILS

All mice were bred and housed in specific pathogen-free conditions on a 12-hr light-dark cycle at ambient temperature in accordance with the Institutional Animal Care and Use Guidelines of the University of North Carolina, Chapel Hill. CD45.1 and CD45.2 C57BL/6 mice aged between 6 and 8 weeks old were used as recipients for cell transfer experiments. Mice bearing a transgenic TCR (P14) recognizing the LCMV GP_33-41_ epitope were used as donor cells throughout the outlined experiments (obtained by Dr. Ananda Goldrath at UCSD). Obtained, generated, and recipient mice were bred in-house, fed standard chow (Purina), and donor and recipient mice were either sex-matched or female cells were transferred into male mice.

## METHOD DETAILS

### T cell transfer, LCMV Infection, and Antibody Treatment

CD45.1 recipient mice received 2 x 10^4^ naïve P14 cells one day prior to intraperitoneal (i.p.) infection with 2 x 10^5^ PFU LCMV Armstrong. Intravascular (i.v.) injection of 5 µg CD45.2 antibody for i.v. labeling of donor cells at 48 and 24 hrs prior to euthanization used either CD45.2 BV510 (Biolegend, clone 104) or CD45.2 AF700 (Biolegend, clone 104). Intravascular injection of 2 µg CD8β antibody for i.v. labeling of donor cells at 3-5 mins prior to euthanization used CD8β AF594 (Biolegend, clone YTS156.7.7).

### Tissue Processing and Spectral Flow Cytometry

Single-cell suspensions from spleens were obtained through mechanical disruption. RBCs were lysed with ACK buffer (150 mM NH4Cl, 1mM KHCO3, and 0.1 mM EDTA, pH 7.4). Intraepithelial lymphocytes were obtained from small intestines through chemical digestion in a buffer containing 154 µg/mL DTE for 30 min at 37°C following removal of Peyer’s patches. Following digestion, single-cell suspensions of lymphocytes were obtained from the small intestine epithelium by filtering cells through a 70-µm nylon filter followed by density gradient spin isolation using 44/67% Cytiva Percoll Centrifugation Media (Fisher Scientific, cat# 45-001-747) for 20 min at 568g.

All samples are stained with fixable live/dead dye (Invitrogen) in FcR block (S17011E; Biolegend) prior to further staining. Staining was performed for 20min at 4°C. Samples were stained with the following antibodies: PD-1 (Biolegend, clone 29F.1A12), Ly6C (Biolegend, clone HK1.4), CX3CR1 (Biolegend, clone SA011F11), KLRG1 (Invitrogen, clone 2F1), CD127 (Biolegend, clone A7R34), CD62L (Biolegend, clone MEL-14), CD52 (Biolegend, clone S20005A), CD25 (Invitrogen, clone PC61.5), CXCR6 (Biolegend, clone SA051D1), CD69 (Biolegend, clone H1.2F3), CD103 (Biolegend, clone 2E7), CD11b (Biolegend, clone M1/70), CCR9 (Biolegend, clone 9B1), and either Vα2 (Biolegend, clone B20.1) or CD45.2 (Biolegend, clone 104).

### Retroviral Transductions

Retroviral transduction of P14 CD8 T cells were performed as previously described (Green et al., 2025) using packaged MSCV retrovirus supplemented with 8 µg/mL polybrene and 50 µM BME, via spin-fection for 1 hr at 37°C and 568g. MSCV retrovirus was generated using PLAT-E packaging cells treated with TransIT-LT1 transfection reagent mixed with 1.5 µg of vector plasmid and 1 µg pcl-eco helper plasmid (1.5:1 ratio) at a 3:1 ratio of TransIT-LT1to DNA. Retrovirus was harvested 48h and 72h after transfection. The following vectors were purchased from Vector Builder for use in these studies: EV-IRES-ametrine, T-bet-T2A-EGFP, TCF1-T2A-EGFP, and AP4-T2A-EGFP. The following vectors used were generously gifted from Ananda Goldrath (UCSD): *Cd19* shRNA-EYFP RV and *Klf2* shRNA-Ametrine RV.

### OMIQ Analysis

Flow cytometry fcs files were compensated and pre-gated to live donor cells and subgated on iv antibody stains in FlowJo (V10.10.0) prior to loading into OMIQ. Files were subsampled to 250-1000 events per file before Leiden clustering analysis was performed. UMAP analysis used default settings and included the following flow markers: PD-1, Ly6C, CX3CR1, KLRG1, CD62L, CD127, CD52, CD103, and CD25. FlowSOM clustering was performed using default settings, except that kvalues was set to 8 and used the same flow markers as the UMAP analysis. Wishbone trajectory analysis was performed using the same flow markers as the UMAP, with C1 set as the starting cell, and waypoint and K set to 250 and 15, respectively.

For downstream analysis of circulating (Figure 2L) and resident (Figures 5H and 5K) CD8 subsets at 28 days p.i., fcs files were loaded into python using flowkit^68^. Flow datasets were then arcsinh-transformed and converted into `anndata` objects for single-cell analysis using `scanpy` ^69^. Specifically, we performed principal component analysis (PCA) with 11 components, and for Figure 5H we fit *k*-means clusters with *k* = 4.

### Processing of scRNAseq CD8 T cell Time Course Data

The scRNAseq dataset consisting of CD8 T cells profiled across post-infection timepoints^12,13^ was downloaded from Gene Expression Omnibus (GEO), with accession number GSE131847. This dataset profiles the response of P14 CD8 T-cells to acutely-resolved LCMV-Armstrong in the siIEL and spleen at 11 timepoints post-infection, plus the Naive (D0) condition. Preprocessing was performed using `scanpy`, and followed a standard scRNA workflow^70^. Steps included filtering out cells with greater than 10% mitochondrial reads or fewer than 1500 total reads, and subsequently filtering out genes detected in fewer than 2 cells; normalizing total counts per cell to 1e6; and log1p transformation.

Highly variable gene selection, principal component analysis (PCA) dimensionality reduction, *k*-nearest neighbors (kNN) and uniform manifold approximation and projection (UMAP) were performed separately for each iteration (spleen-only, combined, and siIEL-only) of the trajectory model. For each version, we selected the top-3000 most variable genes using the `scanpy` implementation of ^71^, performed PCA using 30 components, and fit k-NN using 15 nearest neighbors.

### Cell State Clustering and Annotation

To identify timepoint-specific supercells, we fit kNN at each timepoint using 15 nearest-neighbors and performed Leiden clustering with resolution parameter set to 1. These supercells were then used in Sankey diagrams for gene expression and gene set scores (e.g. Figures 1J, K, and N), visualizing how heterogeneity in CD8 T cell states evolves over time.

To identify the canonical circulating T cell states in Figure 2B, we employed a subsetting approach based on gene set scores. We first obtained gene lists associated with bulk RNA expression of known CD8 subtypes from publicly available datasets GSE157072 ^50^ and GSE173512 ^49^. These datasets consist of bulk RNA expression from flow-sorted CD8 T-cells at timepoints roughly corresponding to those in the scRNA CD8 Infection Timecourse Dataset. We constructed pairwise gene lists of the form l^*A/B*^ by performing differential expression analysis for group *A* vs. group *B*, and selecting the 100 genes with the highest log2-FCs among those with padj < 0.05. We obtained scores *s*^*A/B*^ via the methodology in `sc.tl.score_genes`, as described in ^71^. We then obtained composite scores 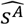 as follows:

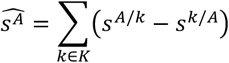

where *K* is the set of all groups excluding *A*. Finally, cells were assigned to the group with the highest score 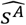. We further imposed prior knowledge-informed restrictions to obtain more biologically-appropriate subsets, namely, we masked scores for the Naïve subset at days p.i. > 5, for the EEC, MP, and TE subsets at days p.i. > 10, and for the TCM, TEM, and t-TEM subsets at days < 14. These restrictions correspond to the windows during which the respective subset terminologies are applied in the literature ^49,50^. This methodology enabled us to identify subsets based on gene set scores that take into consideration both upregulated as well as downregulated genes.

### Optimal Transport

We utilized an entropic unbalanced optimal transport (UOT) framework to fit single-cell differentiation trajectories of CD8 T cells. This is a flexible and robust extension of the Kantorovich formulation ^72^ of OT. We can conceptualize OT in the context of biological trajectories as assigning a discrete joint probability *P*^∗^ over *m* × *n* paths between empirical distributions of *m* cells at a source timepoint to *n* cells at a target timepoint, where the element 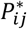 tells us the probability of observing a transition from source cell *i* to target cell *j*. The OT solution *P*^∗^, referred to as a transport map, is defined as,

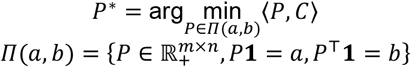

Here, 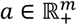 is the probability distribution over source cells, to be transformed into the target distribution, 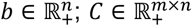 is the cost matrix such that *C*_*ij*_ is the cost of moving a unit probability mass between *a*_*i*_ and *b*_*j*_, defined here as the Euclidean distance ‖***x***_***i***_ − ***x***_***j***_‖_2_ between rows of the cell-by-feature matrix *X* ∈ ℝ^(*m*+*n*)×*d*^; and, *P* ∈ *Π*(*a, b*) is a coupling matrix between source and target distributions within the set of allowable solutions *Π*(*a, b*). Here, we have used angle brackets to denote the Frobenius product as

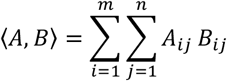

The solution to this objective provides an exact optimal transport solution, however is computationally intensive to obtain and is sensitive to small changes in input data. The addition of a negative entropy term (−*ϵ*ℋ(*P*)) addresses both of these problems by smoothing the loss landscape as,

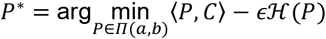

Here, ℋ(*P*) = −⟨*P*,log*P*⟩ is the Shannon entropy of the coupling *P*, and *ϵ* ∈ ℝ_+_ is the magnitude of entropic regularization. As *ϵ* increases, the optimal transport solution gets smoother and less exact. In fact, this smoothness actually makes the solution empirically more robust to noise in the data. Smoothing the loss landscape allows the solution to entropic OT to be found using dual coordinate ascent via the Sinkhorn algorithm ^73^, offering substantial efficiency gains over exact OT.

Unbalanced optimal transport (UOT) relaxes this framework even further, allowing the optimal coupling marginals to deviate from the source and target distributions. This effectively moves the solution away from outliers in the data distribution. The degree of deviation is controlled by hyperparameters λ_*a*_, λ_*b*_ ∈ ℝ_+_, which weights the divergence between the posterior marginals of the OT coupling and the prior marginals of the source and target distributions as,

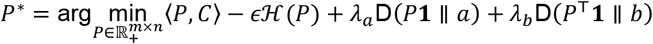

Generally, the Kullback-Liebler (KL) divergence D_KL_(*p* ∥ *q*) = ∑*p*_*i*_log(*p*_*i*_*/q*_*i*_) is used to measure the discrepancy between prior and posterior marginals. With entropic regularization, UOT can be solved using an extension of the efficient Sinkhorn algorithm used to solve balanced entropic OT. The ability to dynamically adjust marginals is particularly useful for single-cell data, where uncertainty in true cell proliferation and apoptosis rates can profoundly impact cell trajectories.

Given *T* timepoints, we fit *T* − 1 optimal transport maps between adjacent timepoints using the `moscot` package ^26^. Trajectory inference in `moscot` involves three main hyperparameters: *ϵ* ∈ ℝ_+_, and *τ*_*a*_, *τ*_*b*_ ∈ [0,1). These parameters control the strength of entropic regularization, and divergence from prior source and target marginals, respectively. The hyperparameters *τ*_*a*_, *τ*_*b*_ are both given by the equation *τ*_*a*_ = λ_*a*_*/*(λ_*a*_ + *ϵ*). For both spleen and siIEL-specific trajectories, we set *ϵ* = 0.01, *τ*_*a*_ = 0.95, *τ*_*b*_ = 0.9995, with *τ*_*b*_ > *τ*_*a*_ implying stronger priors on the target marginals than the source marginals. This accounts for uncertainty in the cell proliferation and death rates. For mixed-compartment trajectories, we set *ϵ* = 0.05, *τ*_*a*_ = 0.95, *τ*_*b*_ = 0.95. Both *ϵ* and *τ*_*b*_ were relaxed to reflect higher uncertainty in the mixed-compartment model, both with respect to cell fate decisions as well as the relative sizes of compartment populations.

For each of the three trajectory models (spleen-only, combined, and siIEL-only), cost matrices were calculated between each pair of adjacent timepoints based on a 50-component local PCA containing only cells from the relevant pair of timepoints, and inheriting the top-3000 highly variable genes from the parent dataset. Cell proliferation and apoptosis scores were calculated using the default gene lists in `moscot.problems.TemporalProblem.score_genes_for_marginals`, and source marginals were estimated using an exponential scaling factor, as described below.

### Tuning Source Marginals

In order to fit biologically realistic trajectories, we adapted methods previously developed in ^24,26^, and refined them to finetune plausible rates of CD8 expansion. Schiebinger et al. proposed using gene set scores associated with cell proliferation and apoptosis signatures to estimate growth rates using a birth-death model, with birth and death rates estimated using a sigmoid function on the proliferation and apoptosis scores, respectively. Klein et al. proposed a simpler parameterization as,

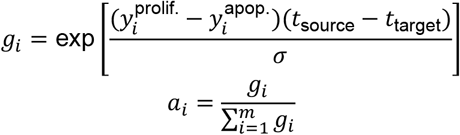

where *g*_*i*_ are the the unnormalized growth rates, *a*_*I*_ are the source marginals, 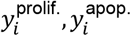 are the proliferation and apoptosis gene set scores, *t*_source_, *t*_target_ are the source and target timepoints, and *σ* is the sole tunable scaling factor.

To prevent large jumps between timepoints, which exaggerate outliers in the proliferation and apoptosis scores and collapse the entire distribution of source marginals to a single cell, we clipped the gene set score difference 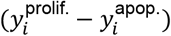 at the 95th percentile within each timepoint to avoid this marginal collapse. Furthermore, we selected the scaling factor *σ* to correspond to the maximum number of cell divisions expected over a 24hr. window. We chose a conservative estimate of 3 cell divisions / day ( 8 hr / division) (De Boer et al., 2001), corresponding to a 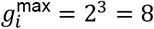. Then, for each timepoint we solved

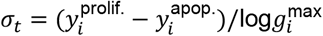

Finally, we set *σ* = max{*σ*_*t*_}. This procedure allowed us to create realistic trajectories of CD8 T cell clonal expansion.

In order to estimate population dynamics, we cached the prior growth rates *g*_*i*_ after fitting OT. Then, we chose the starting population size *N*_0_ = 1000 and calculated population sizes through the recurrence relation:

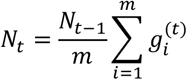

Here, 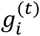 are the growth rates at the timepoint with index *t*. We were further able to find subset population sizes by weighting the total population size at a timepoint *N*_*t*_ by subset proportions.

### Fate Flow Sankey Diagram

To show how cells evolve over time, we used a Sankey diagram that summarizes the probability flow between clusters. For *n* cells over a set of *k* clusters given by the binary matrix *C*^*s*^ ∈ {0,1}^*n*×*k*^ at timepoint *s*, we calculate inflows and outflows separately to account for cell proliferation and death. Given sequential timepoints *r, s, t*: *r* < *s* < *t*, we define the inflows and outflows 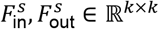 as,

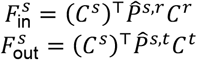

### Quality Controls

To ensure that there were no systematic errors or biases in our OT analysis, we performed a few quality controls for each fitted model. First, we assessed whether unbalanced OT caused substantial deviation between prior and posterior growth estimates. While some differences are expected due to the construction of unbalanced OT, it is desirable to ensure that these are localized, explainable, and unlikely to bias the overall transport map. Next, we evaluated the per-cell source and target OT transport costs, ensuring that there were no cells with extreme influence over the rest of the OT mapping. In general, all transport costs were within an order of magnitude (Figures S1C, S2B, S3C), indicating high-quality mappings. Finally, we ensured that cell cycle phase did not dominate trajectories (Figures S1D, S2C, S3C), indicating that phenotypic state, rather than proliferative status, primarily influenced the structure of the OT trajectories.

### Barycentric Projection and Feature Correlation

To calculate feature correlations over the course of the trajectory, we computed the barycentric projection 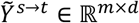 of features from a given timepoint *s* to a timepoint *t* as,

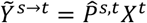

Here, 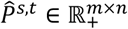 is the row-stochastic OT coupling between timepoints *s, t*, and *X*^*t*^ ∈ ℝ^*n*×*d*^ is the feature matrix at timepoint *t*. We normalize 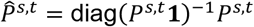 so that every row sums to 1, therefore 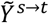 represents the expected feature values at time point *t* under the OT coupling, given cells starting at timepoint *s*.

We computed barycentric projections at all adjacent pairs of timepoints using highly variable genes in both the spleen and siIEL. Then, for the *m* cells at timepoint *s*, we computed the spearman correlation between genes *X*^*s*^ ∈ ℝ^*m*×*d*^ and their barycentric projections 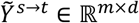, yielding *T* vectors of genewise correlations *ρ*^*t*^ ∈ ℝ^*d*^. We computed the median gene correlations 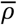 across timepoints, and selected the 100 genes with the highest median correlations 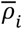 for clustering.

### Trajectory Integration and Clustering

In order to identify salient post-infection trajectories, we developed a methodology to integrate cells from different timepoints based on OT couplings. Our goal was to project disjoint cell distributions from different timepoints into a shared embedding space that smoothly interpolates between temporal cell distributions. We can construct an embedding 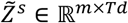 of the entire trajectory of cells across the T timepoints for a timepoint *s* by computing the barycentric projections to all other timepoints 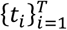, and concatenating them along the feature axis as,

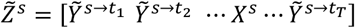

where we have defined 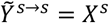. When *s* > *t*, we use *P*^*s,t*^ = (*P*^*t,s*^)^⊤^. For non-adjacent timepoints, we assume that differentiation follows a Markov process and chain the transport maps together, i.e. 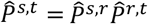 for some *s* < *r* < *t*. This construction allows us to equally weight projections across the entirety of the trajectory for downstream applications, such as clustering.

For the feature matrices *X*^*t*^, we use a *d*-dimensional kernel approximation of the cell density in gene expression space. This allows us to retain higher-order information about cell distributions over time, since the barycentric projection induces a kernel mean embedding. For our analysis, we employed the Nystroem algorithm to fit kernel approximations at each timepoint with *d* = 50. The kernel bandwidths were heuristically determined, by taking the median inter-cell Euclidean distance at each timepoint.

We clustered cells into discrete trajectories based on their embedding vectors 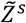, with *s* specifying each cell’s measured timepoint. We employed both leiden and k-means clustering algorithms, and found that both produced quantitatively similar groupings for both the spleen and the siIEL (Figures S1D and S3H). The spleen trajectories were fit using leiden clusters since these were better able to resolve the Naive population; whereas, the siIEL leiden clusters contained several small clusters with only a few cells, so the k-means clusters (which captured the same major groups) were used for downstream analysis. For both compartments, we only clustered on features for timepoints after day 6 p.i. (i.e. 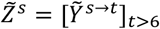), so that the clusters were not confounded by cell migration between compartments.

### Fate Entropy and Consistency Scores

To quantify the uncertainty of cell trajectories, we compute the entropy of cell fate propensities. We define fate propensities 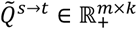 as the probability that a cell at timepoint *s* will transition from one of the m states at timepoint s into one of the k states at the subsequent timepoint, t, as given by the binary cluster matrix accounting for the k possible cluster assignments across the n cells with *C*^*t*^ ∈ {0,1}^*n*×*k*^ as,

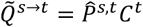

This is equivalent to the barycentric projection of cluster probabilities. Then, we can calculate the Shannon entropy ℋ^*s*→*t*^ ∈ ℝ^*m*^ for each cell:

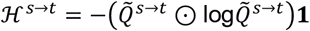

where ⊙ denotes the element-wise (Hadamard) product. In this paper, fate entropy is evaluated for trajectory clusters at the next timepoint.

We extended the fate entropy approach to create a fate consistency score that describes how smoothly a set of clusters divides flows along a trajectory. We chose to consider “smooth” trajectories as those which have minimal amount of uncertainty between timepoints, i.e. those which exhibit low fate entropy. To normalize for differences in cluster size, we compute the expected fate entropy for a set of clusters under a completely random trajectory as,

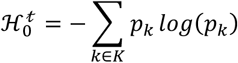

where *K* is the set of clusters and *p*_*k*_ is the frequency of cluster *k* at timepoint *t*. We then define the consistency as 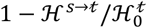. Consistency scores close to 1 indicate clusters that smoothly partition trajectories, while scores below 0 indicate that the clusters give us less information about the trajectory than randomly selecting clusters at the target timepoint.

### Earth Mover’s Distance

We evaluated distributional distances (Figs 2K-L) between populations of cells using optimal transport costs, commonly referred to as the Earth Mover’s Distance (EMD). Specifically, we employed the entropic optimal transport objective, also known as the Sinkhorn distance. Given cell-by-feature matrices *X*^*A*^ ∈ ℝ^*m*×*d*^, *X*^*B*^ ∈ ℝ^*n*×*d*^ for two populations of cells, we calculate the Euclidean distance matrix 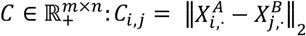. Then, we define the EMD as the cost of the entropic optimal transport solution as

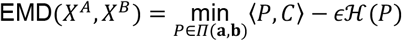

where 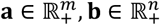 are uniform over the cells in each population. Unlike the exact (un-regularized) optimal transport solution, the Sinkhorn formulation is biased because the entropic regularization ensures that EMD(*X*^*A*^, *X*^*A*^) > 0. We therefore use the unbiased Sinkhorn distance as,

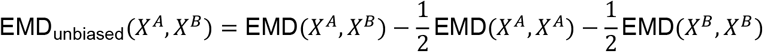

We compute EMDs at each timepoint for cells from a given cluster against cells from all other clusters, using the PCA coordinates and with *ϵ* = 1. For distributions of EMDs, we randomly sample 100 cells from both populations and bootstrap the EMD calculation 10 times.

### Dynamic Time Warping and Gene Clustering

For clustering genes across post-infection timepoints, we used dynamic time warping (DTW) in concert with k-medoids clustering via the dtaidistance Python package (Meert et al. 2020). DTW measures the distances between time series within a sliding window, allowing for smoother time series clusters than clustering on a vector of independent timepoints. Since our post-infection timepoints were coarse, we utilized a sliding window size of 2. For clustering genes in Figures 1 and 5, we averaged the gene expression across the respective compartment at each timepoint, and calculated distances between each resulting time series using DTW.

For clustering regulons in Figure 7, we averaged regulon activity over both spleen and siIEL subsets within each timepoint and concatenated these features into a combined subset/timepoint vector for each regulon. Unlike for gene expression, we did not perform DTW for distance calculation, due to the extra dimension added by subset. Instead, we opted to perform agglomerative hierarchical clustering on the concatenated feature vector.

### SCENIC Pipeline

In order to estimate transcription factor (TF) activity enrichment, we performed gene regulatory network inference using pySCENIC ^63^. The SCENIC algorithm ^62^ infers transcriptional regulatory networks using expression of known TFs to train tree-based models to predict expression for every gene. The resulting models are used to obtain feature importances for each (TF, gene) pair, with each TF representing a “regulon” that controls its target genes. Each regulon is pruned to contain only target genes that contain a TF binding site motif that is both annotated to the TF and above a normalized enrichment score (NES) threshold within a 10 kilo-base pair (kbp) radius of the transcription start site (TSS), or else within 500bp upstream of the TSS. In this analysis, we used an NES threshold of 0.2, and utilized databases of TFs, motif-gene rankings, and motif annotations from https://resources.aertslab.org/cistarget. Finally, we computed activity scores of each regulon in each cell using the AUCell algorithm ^62^. We fit SCENIC regulons on count data for spleen and siIEL across all days simultaneously, using all 9,941 measured genes.

### Differential Gene Expression

We compare gene expression between cell populations using pyDESeq2 ^74^. In brief, this fits a negative-binomial model on gene counts given a design matrix. For comparisons within a single timepoint, we modeled the dependence of gene expression on only the cell subsets. For comparisons across time, we include subset and timepoint as categorical variables, as well as interaction terms between timepoint and subset. In the latter case, we tested hypotheses comparing each subset to the others averaged across time by constructing linear contrasts of the estimated coefficients combining main and interaction effects, weighted by the subset proportions at each timepoint. We then performed a Wald test for the effect size and significance of subset on gene expression. This approach provided a statistically rigorous test of time-averaged differences between cell subsets.

We utilized a similar approach to test for differential activity of SCENIC regulons between subsets, fitting a linear model of regulon activity versus subset and timepoint (categorical) with interactions. We then tested for time averaged effects of subset on regulon activity using the same approach for constructing contrasts as for differential gene expression.

We performed the Benjamini-Hochberg (BH) procedure to control the false discovery rate (FDR). For comparisons of multiple subsets (for example, in Fig. 5), the BH procedure was applied after all tests were performed.

### Experimental statistics

For flow cytometry data analysis, paired or unpaired students t-testing was performed or one-way ANOVA with Tukey’s multiple comparisons as indicated. No outliers were removed from the datasets and mice were randomly assigned to study groups. Symbols are as follows: ns, not-significant; *p≤0.05; **p≤0.01; ***p≤0.005; ****p≤0.001. All data points are shown, bar plots the mean, and error bars are SEM, unless otherwise noted.

---

**Figure S1:**
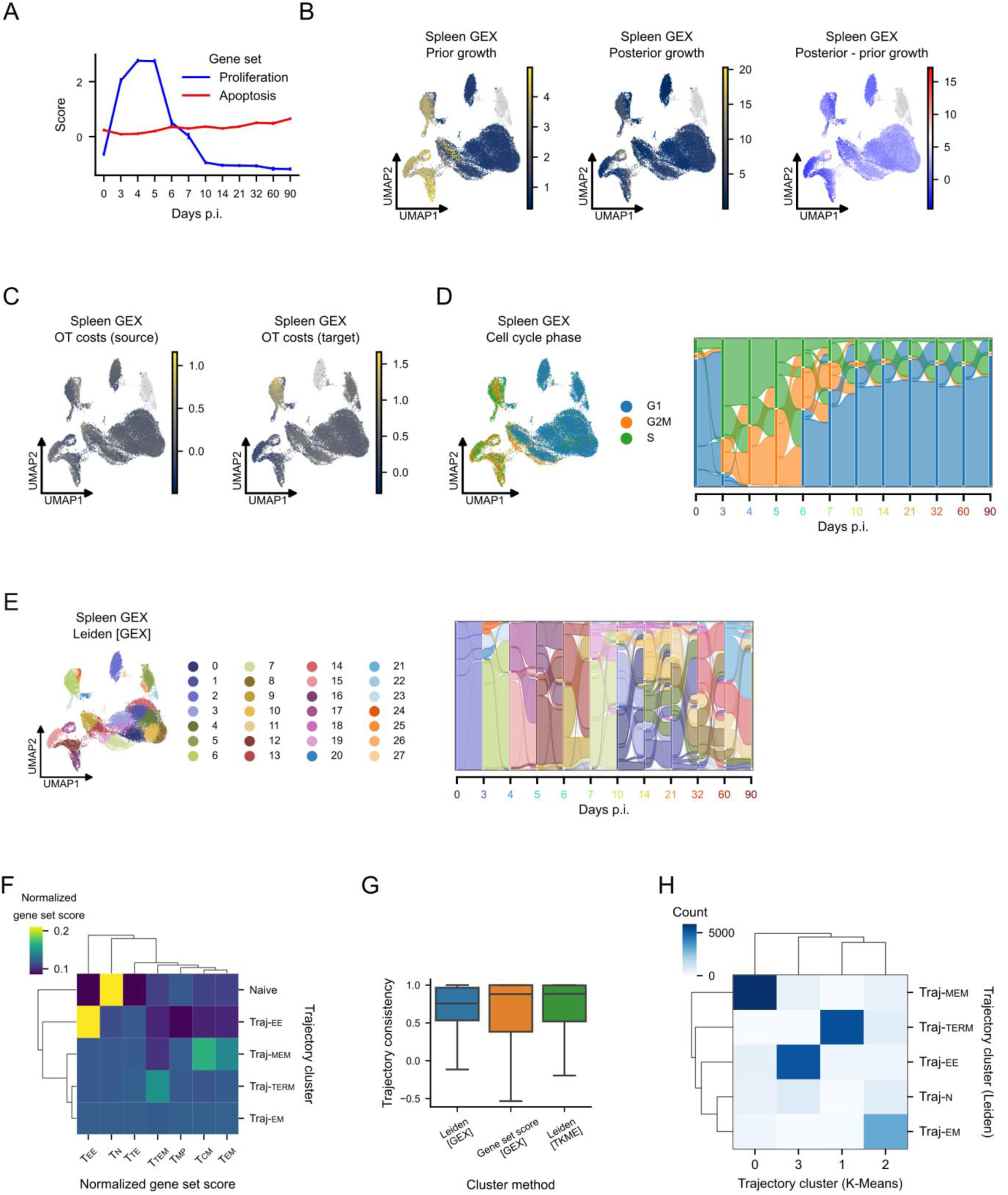
Spleen gene set scores and OT results. **(A)** Mean proliferation (blue) and apoptosis (red) scores over time (error bar = s.e.). **(B)** UMAPs shaded by prior growth rate (left), posterior growth rate (middle), and difference between posterior and prior growth rates (right) used to compute the input OT source marginals corresponding to Figure 1F. **(C)** UMAPs colored by source (left) and target (right) costs of OT trajectory inference in the spleen corresponding to Figure 1F. **(D)** UMAP (left) of estimated cell cycle phase and Sankey diagram (right) showing probability flow of cells from different phases under OT mapping corresponding to Figure 1F. **(E)** UMAP (left) and Sankey diagram (right) of Leiden clusters based on GEX alone corresponding to Figure 2B,G. **(F)** Heatmap of mean gene set scores per trajectory cluster, with each score associated with a canonical cell state. **(G)** Evaluation of clustering methods on gene expression (GEX) and trajectory embeddings using trajectory consistency metric. Values close to 1 are optimal. **(H)** Confusion matrix displaying the co-occurrence of trajectory clusters using different clustering methods.

**Figure S2:**
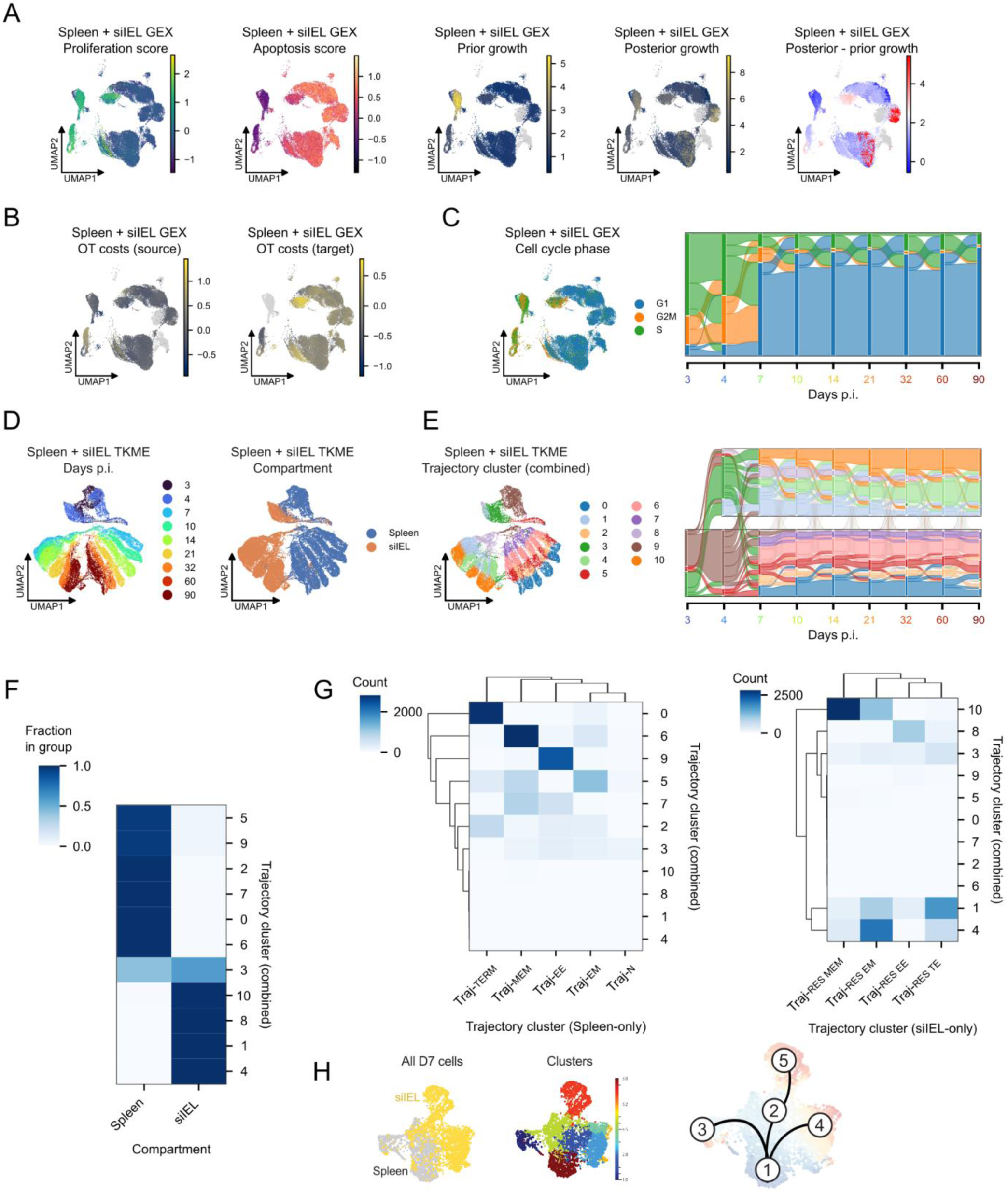
Combined spleen and siIEL gene set scores and OT results. **(A)** UMAPs displaying quantities related to source marginal estimation corresponding to Figure 3A. From left to right: proliferation and apoptosis gene set scores, initial (prior) and posterior estimates of single cell growth rates, and difference between posterior and prior growth rates. **(B)** UMAPs colored by source and target costs of simultaneous OT trajectory inference for both spleen and siIEL. **(C)** UMAP (left) of estimated cell cycle phase and Sankey diagram (right) showing probability flow of cells from different phases under OT mapping. **(D)** UMAP dimensionality-reduction based on TKME features derived from the multi-compartment OT model, colored by day (left) and compartment (right) corresponding to Figure 4A. **(E)** TKME UMAP (left) and Sankey diagram (right) displaying clusters derived from multi-compartment TKME features. **(F)** Confusion matrix displaying fraction of trajectory clusters belonging to each compartment corresponding to Figure 4A. **(G)** Confusion matrices displaying overlap sizes between trajectory clusters from the combined multi-compartment model (rows), and trajectory clusters derived from the spleen-specific (left) and siIEL-specific (right) models (corresponding to Figure 4A). **(H)** Leiden clustering of spectral flow cytometry data from Figure 4G.

**Figure S3:**
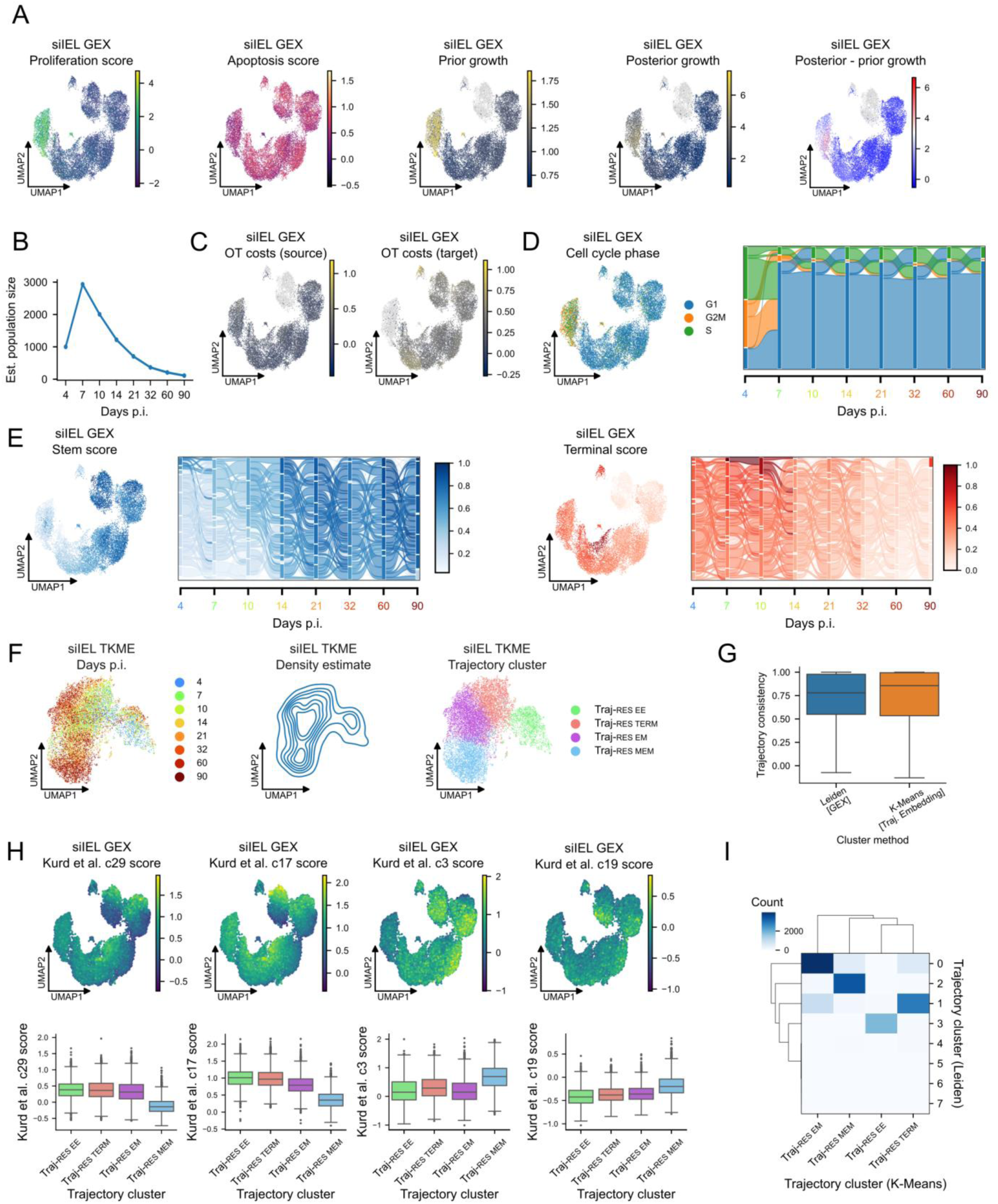
siIEL gene set scores and OT results. **(A)** UMAPs displaying quantities related to source marginal estimation corresponding to Figure 5A. From left to right: proliferation and apoptosis gene set scores, initial (prior) and posterior estimates of single cell growth rates, and difference between posterior and prior growth rates. **(B)** Estimated siIEL-resident CD8 population size, reconstructed from OT model. **(C)** UMAPs shaded by source (left) and target (right) costs of OT trajectory inference in the siIEL. **(D)** UMAP (left) of estimated cell cycle phase and Sankey diagram (right) showing fate flow of cells from different phases under OT mapping. **(E)** UMAPs and Sankey diagrams displaying gene set scores for stemness (left) and terminal differentiation (right) corresponding to Figure 5C. **(F)** UMAP dimensionality-reduction based on TKME features derived from the siIEL-specific OT model, shaded by day (left) and trajectory cluster (right), along with kernel density estimate of cell density in the UMAP space (middle) corresponding to Figure 5D. **(G)** Evaluation of clustering methods on gene expression (GEX) and trajectory embeddings using trajectory consistency metric. Values close to 1 are optimal. **(H)** UMAPs (top) displaying gene set scores for Trm cluster signatures reported in (Kurd et al., 2020)). Boxplots (bottom) of median and IQR gene set scores for each trajectory cluster. **(I)** Confusion matrix displaying the overlap size of trajectory clusters using different clustering methods.

**Figure S4:**
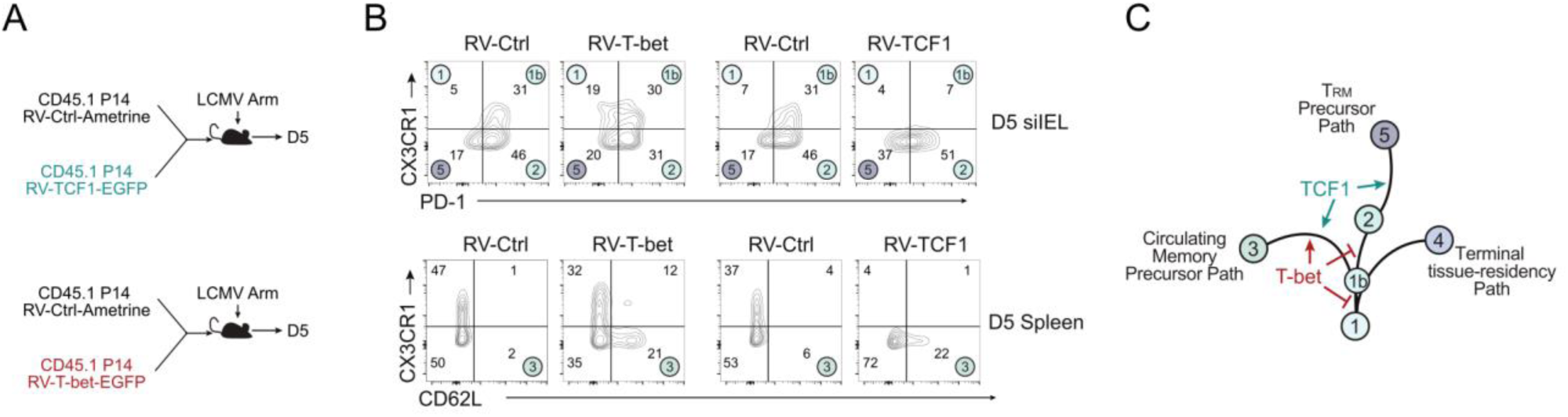
T-bet and TCF1 modulate early circulating and resident differentiation paths. **(A)** Congenic P14 cells transduced with control retrovirus (RV-Ctrl), TCF1-enoding retrovirus (RV-TCF), or T-bet-encoding retrovirus (RV-T-bet) were transferred into mice infected with LCMV Arm and profiled using spectral flow cytometry at D5. **(B)** Flow cytometry analysis of phenotypes in the siIEL (top) and Spleen (bottom). Quadrants are labeled with phenotypes corresponding to SFC clusters from Figure 4M. **(C)** Pseudotime differentiation path from Figure 4M, updated with putative regulatory roles for T-bet and TCF1.

## REFERENCES

1. Chang, J.T., Wherry, E.J., and Goldrath, A.W. (2014). Molecular regulation of effector and memory T cell differentiation. Nat. Immunol. 15, 1104–1115. 10.1038/ni.3031.

2. Koh, C.-H., Lee, S., Kwak, M., Kim, B.-S., and Chung, Y. (2023). CD8 T-cell subsets: heterogeneity, functions, and therapeutic potential. Exp. Mol. Med. 55, 2287–2299. 10.1038/s12276-023-01105-x.

3. Adams, N.M., Grassmann, S., and Sun, J.C. (2020). Clonal expansion of innate and adaptive lymphocytes. Nat. Rev. Immunol. 20, 694–707. 10.1038/s41577-020-0307-4.

4. Masopust, D., and Schenkel, J.M. (2013). The integration of T cell migration, differentiation and function. Nat. Rev. Immunol. 13, 309–320. 10.1038/nri3442.

5. Lavenant, H., Zhang, S., Kim, Y.-H., and Schiebinger, G. (2021). Towards a mathematical theory of trajectory inference. arXiv. 10.48550/arxiv.2102.09204.

6. Huang, H., Zhou, P., Wei, J., Long, L., Shi, H., Dhungana, Y., Chapman, N.M., Fu, G., Saravia, J., Raynor, J.L., et al. (2021). In vivo CRISPR screening reveals nutrient signaling processes underpinning CD8+ T cell fate decisions. Cell 184, 1245–1261.e21. 10.1016/j.cell.2021.02.021.

7. Yoon, H., Kim, T.S., and Braciale, T.J. (2010). The cell cycle time of CD8+ T cells responding in vivo is controlled by the type of antigenic stimulus. PLoS ONE 5, e15423. 10.1371/journal.pone.0015423.

8. He, S., Zhu, Y., Tavakol, D.N., Ye, H., Lao, Y.-H., Zhu, Z., Xu, C., Chauhan, S., Garty, G., Tomer, R., et al. (2026). Squidiff: predicting cellular development and responses to perturbations using a diffusion model. Nat. Methods 23, 65–77. 10.1038/s41592-025-02877-y.

9. Park, C., Mani, S., Beltran-Velez, N., Maurer, K., Huang, T., Li, S., Gohil, S., Livak, K.J., Knowles, D.A., Wu, C.J., et al. (2024). A Bayesian framework for inferring dynamic intercellular interactions from time-series single-cell data. Genome Res. 34, 1384–1396. 10.1101/gr.279126.124.

10. Bailey, S.R., Bartee, E., Daniels, K.G., Heery, C.R., Kaumaya, P., Lesinski, G.B., Lowinger, T.B., Nelson, M.H., Rubinstein, M.P., Wittling, M.C., et al. (2025). Constructing the cure: engineering the next wave of antibody and cellular immune therapies. J. Immunother. Cancer 13. 10.1136/jitc-2025-011761.

11. Green, W.D., Gomez, A., Plotkin, A.L., Pratt, B.M., Merritt, E.F., Mullins, G.N., Kren, N.P., Modliszewski, J.L., Zhabotynsky, V., Woodcock, M.G., et al. (2025). Enhancer-driven gene regulatory networks reveal transcription factors governing T cell adaptation and differentiation in the tumor microenvironment. Immunity 58, 1725–1741.e9. 10.1016/j.immuni.2025.04.030.

12. Milner, J.J., Toma, C., He, Z., Kurd, N.S., Nguyen, Q.P., McDonald, B., Quezada, L., Widjaja, C.E., Witherden, D.A., Crowl, J.T., et al. (2020). Heterogenous Populations of Tissue-Resident CD8+ T Cells Are Generated in Response to Infection and Malignancy. Immunity 52, 808–824.e7. 10.1016/j.immuni.2020.04.007.

13. Kurd, N.S., He, Z., Louis, T.L., Milner, J.J., Omilusik, K.D., Jin, W., Tsai, M.S., Widjaja, C.E., Kanbar, J.N., Olvera, J.G., et al. (2020). Early precursors and molecular determinants of tissue-resident memory CD8+ T lymphocytes revealed by single-cell RNA sequencing. Sci. Immunol. 5. 10.1126/sciimmunol.aaz6894.

14. Chung, H.K., Liu, C., Battu, A., Jambor, A.N., Pratt, B.M., Xie, F., Riesenberg, B.P., Casillas, E., Sun, M., Landoni, E., et al. (2025). Atlas-Guided Discovery of Transcription Factors for T Cell Programming. BioRxiv. 10.1101/2023.01.03.522354.

15. Chu, Y., Dai, E., Li, Y., Han, G., Pei, G., Ingram, D.R., Thakkar, K., Qin, J.-J., Dang, M., Le, X., et al. (2023). Pan-cancer T cell atlas links a cellular stress response state to immunotherapy resistance. Nat. Med. 29, 1550–1562. 10.1038/s41591-023-02371-y.

16. Buquicchio, F.A., Fonseca, R., Yan, P.K., Wang, F., Evrard, M., Obers, A., Gutierrez, J.C., Raposo, C.J., Belk, J.A., Daniel, B., et al. (2024). Distinct epigenomic landscapes underlie tissue-specific memory T cell differentiation. Immunity 57, 2202–2215.e6. 10.1016/j.immuni.2024.06.014.

17. Street, K., Risso, D., Fletcher, R.B., Das, D., Ngai, J., Yosef, N., Purdom, E., and Dudoit, S. (2018). Slingshot: cell lineage and pseudotime inference for single-cell transcriptomics. BMC Genomics 19, 477. 10.1186/s12864-018-4772-0.

18. Haghverdi, L., Büttner, M., Wolf, F.A., Buettner, F., and Theis, F.J. (2016). Diffusion pseudotime robustly reconstructs lineage branching. Nat. Methods 13, 845–848. 10.1038/nmeth.3971.

19. Bergen, V., Lange, M., Peidli, S., Wolf, F.A., and Theis, F.J. (2020). Generalizing RNA velocity to transient cell states through dynamical modeling. Nat. Biotechnol. 38, 1408–1414. 10.1038/s41587-020-0591-3.

20. Hudson, W.H., and Wieland, A. (2023). Technology meets TILs: Deciphering T cell function in the-omics era. Cancer Cell 41, 41–57. 10.1016/j.ccell.2022.09.011.

21. Youngblood, B., Hale, J.S., Kissick, H.T., Ahn, E., Xu, X., Wieland, A., Araki, K., West, E.E., Ghoneim, H.E., Fan, Y., et al. (2017). Effector CD8 T cells dedifferentiate into long-lived memory cells. Nature 552, 404–409. 10.1038/nature25144.

22. Herndler-Brandstetter, D., Ishigame, H., Shinnakasu, R., Plajer, V., Stecher, C., Zhao, J., Lietzenmayer, M., Kroehling, L., Takumi, A., Kometani, K., et al. (2018). KLRG1+ Effector CD8+ T Cells Lose KLRG1, Differentiate into All Memory T Cell Lineages, and Convey Enhanced Protective Immunity. Immunity 48, 716–729.e8. 10.1016/j.immuni.2018.03.015.

23. Masopust, D., Choo, D., Vezys, V., Wherry, E.J., Duraiswamy, J., Akondy, R., Wang, J., Casey, K.A., Barber, D.L., Kawamura, K.S., et al. (2010). Dynamic T cell migration program provides resident memory within intestinal epithelium. J. Exp. Med. 207, 553–564. 10.1084/jem.20090858.

24. Schiebinger, G., Shu, J., Tabaka, M., Cleary, B., Subramanian, V., Solomon, A., Gould, J., Liu, S., Lin, S., Berube, P., et al. (2019). Optimal-Transport Analysis of Single-Cell Gene Expression Identifies Developmental Trajectories in Reprogramming. Cell 176, 928–943.e22. 10.1016/j.cell.2019.01.006.

25. Bunne, C., Meng-Papaxanthos, L., Krause, A., and Cuturi, M. (2021). Proximal Optimal Transport Modeling of Population Dynamics. arXiv. 10.48550/arxiv.2106.06345.

26. Klein, D., Palla, G., Lange, M., Klein, M., Piran, Z., Gander, M., Meng-Papaxanthos, L., Sterr, M., Saber, L., Jing, C., et al. (2025). Mapping cells through time and space with moscot. Nature 638, 1065–1075. 10.1038/s41586-024-08453-2.

27. Gorin, G., Fang, M., Chari, T., and Pachter, L. (2022). RNA velocity unraveled. PLoS Comput. Biol. 18, e1010492. 10.1371/journal.pcbi.1010492.

28. Reina-Campos, M., Monell, A., Ferry, A., Luna, V., Cheung, K.P., Galletti, G., Scharping, N.E., Takehara, K.K., Quon, S., Challita, P.P., et al. (2025). Tissue-resident memory CD8 T cell diversity is spatiotemporally imprinted. Nature 639, 483–492. 10.1038/s41586-024-08466-x.

29. Gill, A.L., Hudson, W.H., Valanparambil, R.M., Ahn, E., McGuire, D.J., Wieland, A., McManus, D.T., Kissick, H.T., Akondy, R.S., and Ahmed, R. (2023). Longitudinal Analysis of the Phenotype, Transcriptional Profile, and Anatomic Location of Memory CD8 T Cell Subsets after Acute Viral Infection. J. Virol. 97, e0155622. 10.1128/jvi.01556-22.

30. Zhang, N., and Bevan, M.J. (2011). CD8(+) T cells: foot soldiers of the immune system. Immunity 35, 161–168. 10.1016/j.immuni.2011.07.010.

31. Giles, J.R., Ngiow, S.F., Manne, S., Baxter, A.E., Khan, O., Wang, P., Staupe, R., Abdel-Hakeem, M.S., Huang, H., Mathew, D., et al. (2022). Shared and distinct biological circuits in effector, memory and exhausted CD8+ T cells revealed by temporal single-cell transcriptomics and epigenetics. Nat. Immunol. 23, 1600–1613. 10.1038/s41590-022-01338-4.

32. Kaech, S.M., and Cui, W. (2012). Transcriptional control of effector and memory CD8+ T cell differentiation. Nat. Rev. Immunol. 12, 749–761. 10.1038/nri3307.

33. Gerlach, C., Rohr, J.C., Perié, L., van Rooij, N., van Heijst, J.W.J., Velds, A., Urbanus, J., Naik, S.H., Jacobs, H., Beltman, J.B., et al. (2013). Heterogeneous differentiation patterns of individual CD8+ T cells. Science 340, 635–639. 10.1126/science.1235487.

34. Mold, J.E., Modolo, L., Hård, J., Zamboni, M., Larsson, A.J.M., Stenudd, M., Eriksson, C.-J., Durif, G., Ståhl, P.L., Borgström, E., et al. (2021). Divergent clonal differentiation trajectories establish CD8+ memory T cell heterogeneity during acute viral infections in humans. Cell Rep. 35, 109174. 10.1016/j.celrep.2021.109174.

35. Mold, J.E., Weissman, M.H., Ratz, M., Hagemann-Jensen, M., Hård, J., Eriksson, C.-J., Toosi, H., Berghenstråhle, J., Ziegenhain, C., von Berlin, L., et al. (2024). Clonally heritable gene expression imparts a layer of diversity within cell types. Cell Syst. 15, 149–165.e10. 10.1016/j.cels.2024.01.004.

36. Roychoudhuri, R., Lefebvre, F., Honda, M., Pan, L., Ji, Y., Klebanoff, C.A., Nichols, C.N., Fourati, S., Hegazy, A.N., Goulet, J.-P., et al. (2015). Transcriptional profiles reveal a stepwise developmental program of memory CD8(+) T cell differentiation. Vaccine 33, 914–923. 10.1016/j.vaccine.2014.10.007.

37. Daniel, B., Yost, K.E., Hsiung, S., Sandor, K., Xia, Y., Qi, Y., Hiam-Galvez, K.J., Black, M., J Raposo, C., Shi, Q., et al. (2022). Divergent clonal differentiation trajectories of T cell exhaustion. Nat. Immunol. 23, 1614–1627. 10.1038/s41590-022-01337-5.

38. Quezada, L.K., Jin, W., Liu, Y.C., Kim, E.S., He, Z., Indralingam, C.S., Tysl, T., Labarta-Bajo, L., Wehrens, E.J., Jo, Y., et al. (2023). Early transcriptional and epigenetic divergence of CD8+ T cells responding to acute versus chronic infection. PLoS Biol. 21, e3001983. 10.1371/journal.pbio.3001983.

39. Gearty, S.V., Dündar, F., Zumbo, P., Espinosa-Carrasco, G., Shakiba, M., Sanchez-Rivera, F.J., Socci, N.D., Trivedi, P., Lowe, S.W., Lauer, P., et al. (2022). An autoimmune stem-like CD8 T cell population drives type 1 diabetes. Nature 602, 156–161. 10.1038/s41586-021-04248-x.

40. Ellis, G.I., Sheppard, N.C., and Riley, J.L. (2021). Genetic engineering of T cells for immunotherapy. Nat. Rev. Genet. 22, 427–447. 10.1038/s41576-021-00329-9.

41. Pariset, M., Hsieh, Y.-P., Bunne, C., Krause, A., and De Bortoli, V. (2023). Unbalanced Diffusion Schrödinger Bridge. arXiv. 10.48550/arxiv.2306.09099.

42. Kaech, S.M., and Wherry, E.J. (2007). Heterogeneity and cell-fate decisions in effector and memory CD8+ T cell differentiation during viral infection. Immunity 27, 393–405. 10.1016/j.immuni.2007.08.007.

43. Abadie, K., Clark, E.C., Valanparambil, R.M., Ukogu, O., Yang, W., Daza, R.M., Ng, K.K.H., Fathima, J., Wang, A.L., Lee, J., et al. (2024). Reversible, tunable epigenetic silencing of TCF1 generates flexibility in the T cell memory decision. Immunity 57, 271–286.e13. 10.1016/j.immuni.2023.12.006.

44. Althaus, C.L., Ganusov, V.V., and De Boer, R.J. (2007). Dynamics of CD8+ T cell responses during acute and chronic lymphocytic choriomeningitis virus infection. J. Immunol. 179, 2944–2951. 10.4049/jimmunol.179.5.2944.

45. De Boer, R.J., Oprea, M., Antia, R., Murali-Krishna, K., Ahmed, R., and Perelson, A.S. (2001). Recruitment times, proliferation, and apoptosis rates during the CD8(+) T-cell response to lymphocytic choriomeningitis virus. J. Virol. 75, 10663–10669. 10.1128/JVI.75.22.10663-10669.2001.

46. Wu, H., Kumar, A., Miao, H., Holden-Wiltse, J., Mosmann, T.R., Livingstone, A.M., Belz, G.T., Perelson, A.S., Zand, M.S., and Topham, D.J. (2011). Modeling of influenza-specific CD8+ T cells during the primary response indicates that the spleen is a major source of effectors. J. Immunol. 187, 4474–4482. 10.4049/jimmunol.1101443.

47. McInnes, L., Healy, J., and Melville, J. (2020). UMAP: Uniform Manifold Approximation and Projection for Dimension Reduction. arXiv:1802.03426 [cs, stat].

48. Wherry, E.J., Blattman, J.N., Murali-Krishna, K., van der Most, R., and Ahmed, R. (2003). Viral persistence alters CD8 T-cell immunodominance and tissue distribution and results in distinct stages of functional impairment. J. Virol. 77, 4911–4927. 10.1128/jvi.77.8.4911-4927.2003.

49. Milner, J.J., Toma, C., Quon, S., Omilusik, K., Scharping, N.E., Dey, A., Reina-Campos, M., Nguyen, H., Getzler, A.J., Diao, H., et al. (2021). Bromodomain protein BRD4 directs and sustains CD8 T cell differentiation during infection. J. Exp. Med. 218. 10.1084/jem.20202512.

50. Milner, J.J., Nguyen, H., Omilusik, K., Reina-Campos, M., Tsai, M., Toma, C., Delpoux, A., Boland, B.S., Hedrick, S.M., Chang, J.T., et al. (2020). Delineation of a molecularly distinct terminally differentiated memory CD8 T cell population. Proc Natl Acad Sci USA 117, 25667–25678. 10.1073/pnas.2008571117.

51. Renkema, K.R., Huggins, M.A., Borges da Silva, H., Knutson, T.P., Henzler, C.M., and Hamilton, S.E. (2020). KLRG1+ Memory CD8 T Cells Combine Properties of Short-Lived Effectors and Long-Lived Memory. J. Immunol. 205, 1059–1069. 10.4049/jimmunol.1901512.

52. Fagerberg, E., Attanasio, J., Dien, C., Singh, J., Kessler, E.A., Abdullah, L., Shen, J., Hunt, B.G., Connolly, K.A., De Brouwer, E., et al. (2025). KLF2 maintains lineage fidelity and suppresses CD8 T cell exhaustion during acute LCMV infection. Science 387, eadn2337. 10.1126/science.adn2337.

53. Carlson, C.M., Endrizzi, B.T., Wu, J., Ding, X., Weinreich, M.A., Walsh, E.R., Wani, M.A., Lingrel, J.B., Hogquist, K.A., and Jameson, S.C. (2006). Kruppel-like factor 2 regulates thymocyte and T-cell migration. Nature 442, 299–302. 10.1038/nature04882.

54. Masopust, D., and Soerens, A.G. (2019). Tissue-Resident T Cells and Other Resident Leukocytes. Annu. Rev. Immunol. 37, 521–546. 10.1146/annurev-immunol-042617-053214.

55. Milner, J.J., Toma, C., Yu, B., Zhang, K., Omilusik, K., Phan, A.T., Wang, D., Getzler, A.J., Nguyen, T., Crotty, S., et al. (2017). Runx3 programs CD8+ T cell residency in non-lymphoid tissues and tumours. Nature 552, 253–257. 10.1038/nature24993.

56. Mackay, L.K., Minnich, M., Kragten, N.A.M., Liao, Y., Nota, B., Seillet, C., Zaid, A., Man, K., Preston, S., Freestone, D., et al. (2016). Hobit and Blimp1 instruct a universal transcriptional program of tissue residency in lymphocytes. Science 352, 459–463. 10.1126/science.aad2035.

57. Shanahan, S.-L., Kunder, N., Inaku, C., Hagan, N.B., Gibbons, G., Mathey-Andrews, N., Anandappa, G., Soares, S., Pauken, K.E., Jacks, T., et al. (2024). Longitudinal intravascular antibody labeling identified regulatory T cell recruitment as a therapeutic target in a mouse model of lung cancer. J. Immunol. 213, 906–918. 10.4049/jimmunol.2400268.

58. Mackay, L.K., Rahimpour, A., Ma, J.Z., Collins, N., Stock, A.T., Hafon, M.-L., Vega-Ramos, J., Lauzurica, P., Mueller, S.N., Stefanovic, T., et al. (2013). The developmental pathway for CD103(+)CD8+ tissue-resident memory T cells of skin. Nat. Immunol. 14, 1294–1301. 10.1038/ni.2744.

59. Sheridan, B.S., Pham, Q.-M., Lee, Y.-T., Cauley, L.S., Puddington, L., and Lefrançois, L. (2014). Oral infection drives a distinct population of intestinal resident memory CD8(+) T cells with enhanced protective function. Immunity 40, 747–757. 10.1016/j.immuni.2014.03.007.

60. Walunas, T.L., Bruce, D.S., Dustin, L., Loh, D.Y., and Bluestone, J.A. (1995). Ly-6C is a marker of memory CD8+ T cells. J. Immunol. 155, 1873–1883.

61. Hänninen, A., Maksimow, M., Alam, C., Morgan, D.J., and Jalkanen, S. (2011). Ly6C supports preferential homing of central memory CD8+ T cells into lymph nodes. Eur. J. Immunol. 41, 634–644. 10.1002/eji.201040760.

62. Aibar, S., González-Blas, C.B., Moerman, T., Huynh-Thu, V.A., Imrichova, H., Hulselmans, G., Rambow, F., Marine, J.-C., Geurts, P., Aerts, J., et al. (2017). SCENIC: single-cell regulatory network inference and clustering. Nat. Methods 14, 1083–1086. 10.1038/nmeth.4463.

63. Van de Sande, B., Flerin, C., Davie, K., De Waegeneer, M., Hulselmans, G., Aibar, S., Seurinck, R., Saelens, W., Cannoodt, R., Rouchon, Q., et al. (2020). A scalable SCENIC workflow for single-cell gene regulatory network analysis. Nat. Protoc. 15, 2247–2276. 10.1038/s41596-020-0336-2.

64. Mackay, L.K., Wynne-Jones, E., Freestone, D., Pellicci, D.G., Mielke, L.A., Newman, D.M., Braun, A., Masson, F., Kallies, A., Belz, G.T., et al. (2015). T-box Transcription Factors Combine with the Cytokines TGF-β and IL-15 to Control Tissue-Resident Memory T Cell Fate. Immunity 43, 1101–1111. 10.1016/j.immuni.2015.11.008.

65. Yu, B., Zhang, K., Milner, J.J., Toma, C., Chen, R., Scott-Browne, J.P., Pereira, R.M., Crotty, S., Chang, J.T., Pipkin, M.E., et al. (2017). Epigenetic landscapes reveal transcription factors that regulate CD8+ T cell differentiation. Nat. Immunol. 18, 573–582. 10.1038/ni.3706.

66. Laidlaw, B.J., Zhang, N., Marshall, H.D., Staron, M.M., Guan, T., Hu, Y., Cauley, L.S., Craft, J., and Kaech, S.M. (2014). CD4+ T cell help guides formation of CD103+ lung-resident memory CD8+ T cells during influenza viral infection. Immunity 41, 633–645. 10.1016/j.immuni.2014.09.007.

67. Chou, C., Verbaro, D.J., Tonc, E., Holmgren, M., Cella, M., Colonna, M., Bhattacharya, D., and Egawa, T. (2016). The Transcription Factor AP4 Mediates Resolution of Chronic Viral Infection through Amplification of Germinal Center B Cell Responses. Immunity 45, 570–582. 10.1016/j.immuni.2016.07.023.

68. White, S., Quinn, J., Enzor, J., Staats, J., Mosier, S.M., Almarode, J., Denny, T.N., Weinhold, K.J., Ferrari, G., and Chan, C. (2021). Flowkit: A python toolkit for integrated manual and automated cytometry analysis workflows. Front. Immunol. 12, 768541. 10.3389/fimmu.2021.768541.

69. Wolf, F.A., Angerer, P., and Theis, F.J. (2018). SCANPY: large-scale single-cell gene expression data analysis. Genome Biol. 19, 15. 10.1186/s13059-017-1382-0.

70. Luecken, M.D., and Theis, F.J. (2019). Current best practices in single-cell RNA-seq analysis: a tutorial. Mol. Syst. Biol. 15, e8746. 10.15252/msb.20188746.

71. Satija, R., Farrell, J.A., Gennert, D., Schier, A.F., and Regev, A. (2015). Spatial reconstruction of single-cell gene expression data. Nat. Biotechnol. 33, 495–502. 10.1038/nbt.3192.

72. Peyré, G., and Cuturi, M. (2019). Computational Optimal Transport. FNT in Machine Learning 11, 355–206. 10.1561/2200000073.

73. Cuturi, M. (2013). Sinkhorn Distances: Lightspeed Computation of Optimal Transport. In Advances in Neural Information Processing Systems (Curran Associates, Inc.).

74. Muzellec, B., Teleńczuk, M., Cabeli, V., and Andreux, M. (2023). PyDESeq2: a python package for bulk RNA-seq differential expression analysis. Bioinformatics 39. 10.1093/bioinformatics/btad547.

